# Genetic modulation of initial sensitivity to Δ9-tetrahydrocannabinol (THC) among the BXD family of mice

**DOI:** 10.1101/2021.01.08.425948

**Authors:** C. Parks, C.M. Rogers, J.P. Prins, R.W. Williams, H. Chen, B.C. Jones, B.M. Moore, M.K. Mulligan

## Abstract

Cannabinoid receptor 1 activation by the major psychoactive component in cannabis, Δ9-tetrahydrocannabinol (THC), produces motor impairments, hypothermia, and analgesia upon acute exposure. In previous work, we demonstrated significant sex and strain differences in acute responses to THC following administration of a single dose (10 mg/kg, *i.p*.) in C57BL/6J (B6) and DBA/2J (D2) inbred mice. To determine the extent to which these differences are heritable, we quantified acute responses to a single dose of THC (10 mg/kg, *i.p*.) in males and females from 20 members of the BXD family of inbred strains derived by crossing and inbreeding B6 and D2 mice. Acute THC responses (initial sensitivity) were quantified as changes from baseline for: 1. spontaneous activity in the open field (mobility), 2. body temperature (hypothermia), and 3. tail withdrawal latency to a thermal stimulus (analgesia/antinociception). Initial sensitivity to the immobilizing, hypothermic, and antinociceptive effects of THC varied substantially across the BXD family. Heritability was highest for mobility and hypothermia traits, indicating that segregating genetic variants modulate initial sensitivity to THC. We identified genomic loci and candidate genes, including *Ndufs2, Scp2, Rps6kb1* or P70S6K, *Pde4d*, and *Pten*, that may control variation in THC initial sensitivity. We also detected strong correlations between initial responses to THC and legacy phenotypes related to intake or response to other drugs of abuse (cocaine, ethanol, and morphine). Our study demonstrates the feasibility of mapping genes and variants modulating THC responses in the BXDs to systematically define biological processes and liabilities associated with drug use and abuse.

## INTRODUCTION

Recombinant inbred (RI) rodent populations are a valuable resource for forward genetic mapping and systems genetics analysis. Individual RI lines are stable and each genotype can be resampled to boost mapping power and improve quantitative trait loci (QTL) detection (Phillips, Belknap, and Crabbe 1991). In addition, RI population genotypes and trait data collected across time and laboratory settings can be combined into a powerful database of legacy phenotypes amenable to systems scale analyses both across and between trait levels. Here, we leverage the BXD RI family of strains originally derived by crossing C57BL/6J (B6) and DBA/2J (D2) mice followed by intercrossing and inbreeding of their progeny to yield stable RI lines (**Figure 1A**). Since its inception in the late 1970’s, the BXD RI panel continues to be a valuable resource for characterization of the genetic architecture of addiction-related behavior and identification of genes and variants modulating the response to drugs of abuse (Dickson et al. 2016; Philip et al. 2010; Hitzemann et al. 2003; Rodriguez et al. 1994, 1995; Phillips, Belknap, and Crabbe 1991; Jackson et al. 2011).

**Figure 1.**
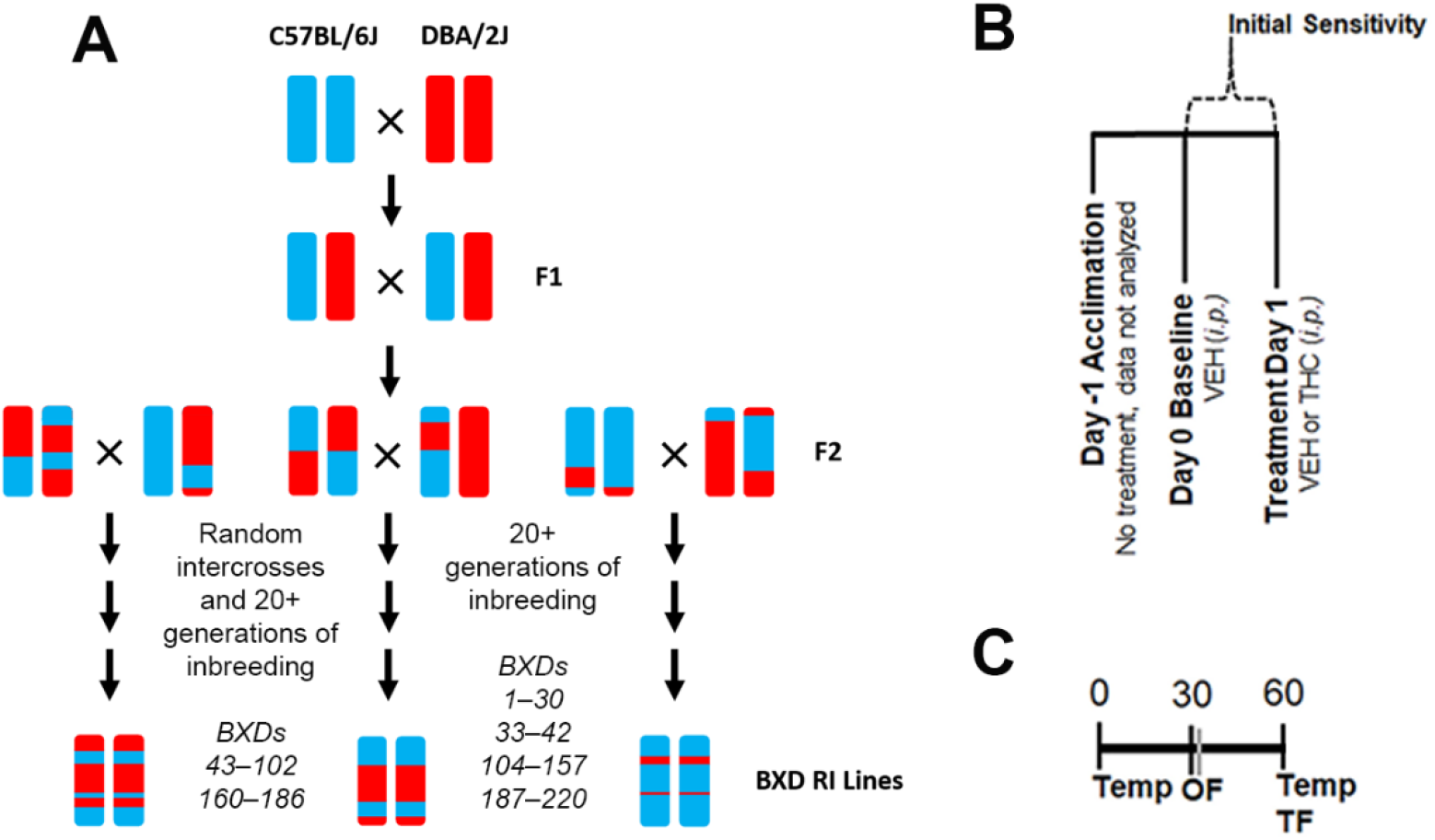
Experimental Overview. (A) Overview of the generation of the BXD RI population. This family of strains has been derived at several points in time from both F2 and advanced intercrosses and there are now ~150 strains available. (B) Description of daily treatment regime. Initial sensitivity is measured by subtracting Day 1 trait values from those of Day 0 for each individual. (C) Timing of daily trait measurements for body temperature (Temp), time mobile in the open field (OF), and latency to withdrawal the tail in response to a thermal stimulus (TF) are shown as minutes post-injection.

With well over 100 BXD lines currently available (Ashbrook et al. 2021), the BXD RI family is the largest and best characterized rodent genetic reference population available (Mulligan et al. 2017; Parker et al. 2017; Ashbrook et al. 2021). This family contains over 6 million segregating variants and ~10,000 recombinations resulting in comparatively high mapping precision (Ashbrook et al. 2021). The BXD family has also been deeply phenotyped for molecular, behavioral, physiological, and pharmacological traits since the late 1970’s and includes data related to cognitive function, anxiety and stress, social interactions, and response to drugs of abuse (Newbury and Rosen 2012; Laughlin et al. 2011; Knoll, Jiang, and Levitt 2018; Philip et al. 2010; Harenza et al. 2014; Loos et al. 2014; Neuner et al. 2016; Ashbrook et al. 2018).

Previously, we established that the parents of the BXDs (strains B6 and D2) differ greatly in their initial responses to a single dose (10 mg/kg, *i.p*.) of THC, the major psychoactive component in cultivars of cannabis (Parks et al. 2020). We have also demonstrated that striatal protein levels of the primary target of THC, the cannabinoid receptor 1 (CB1R), are significantly higher in B6 compared to D2 mice (Parks et al. 2019). However, there are no high impact segregating variants within the CB1R gene locus of B6 and D2 inbred strains that can account for the variation in receptor levels (Parks et al. 2019). Thus, the causal genes and variants that underlie differences in response to THC between B6 and D2 and innate variation in the endogenous cannabinoid receptor signaling pathway remain to be discovered. In this study, we leverage the BXD family to address two main questions:

1. Are differences in THC responses between B6 and D2 inbred strains heritable and, if so, do these differences also segregate among the BXD family?
2. Are initial responses to THC related to responses to other drugs of abuse?

Answers in the affirmative would greatly support the use of the BXD family to identify DNA variants, genes, and shared causal networks mediating responses to THC and other abused substances.

## RESULTS

### Acute THC treatment has a profound and significant effect on mobility, hypothermia, and analgesia initial response traits

Activation of CB1R in discrete neuronal populations (Monory et al. 2007) by THC results in a well characterized trait spectrum that includes a reduction in spontaneous locomotion, hypothermia, and analgesia/antinociception. Genetic or pharmacological deletion of CB1R abrogates these effects (Compton et al. 1996; Huestis et al. 2001; Ledent et al. 1999; Rinaldi-Carmona et al. 1994; Zimmer et al. 1999). Thus, quantification of mobility, hypothermia, and analgesia following THC exposure is a robust method to measure response to THC mediated specifically through CB1R signaling pathways.

We found that treatment (*i.e*. baseline values on day 0 compared to day 1 values following THC treatment) had significant (all *p*-values < 0.001) and moderate to large main effects on mobility, hypothermia, and analgesia in our BXD cohort. Relative to the VEH treatment on day 0, THC treatment on day 1 greatly and significantly reduced mobility (F1,220 = 743.11, p < 0.001, ωp^2^ = 0.71, 90% CI [0.66, 0.75]). THC treatment caused a large and significant decrease in body temperature (F1,232 = 588.27, p < 0.001, ωp^2^ = 0.65, 90% CI [0.60, 0.70]). Acute THC treatment resulted in a moderate and significant increase in tail withdrawal latency (F1,230 = 141.34, p < 0.001, ωp^2^ = 0.31, 90% CI [0.23, 0.38]).

Relative to treatment effects, main effects of strain or sex were generally smaller. Strain effects contributed significantly to response variation for mobility (F21,220 = 7.74, p < 0.001, ωp^2^ = 0.32, 90% CI [0.19, 0.36]), hypothermia (F21,232 = 3.0, p < 0.01, ωp^2^ = 0.12, 90% CI [0.00, 0.11]), and analgesia (F21,230 = 2.89, p < 0.001, ωp^2^ = 0.11, 90% CI [0.00, 0.11]). Significant main effects of sex were observed for hypothermia (F1,232 = 11.07, p < 0.01, ωp^2^ = 0.03, 90% CI [0.00, 0.08]) and analgesia (F1,230 = 7.24, p < 0.01, ωp^2^ = 0.02, 90% CI [0.00, 0.06]).

We observed a moderate and significant treatment-by-strain interaction effect for mobility (F21,220 = 7.40, p < 0.001, ωp^2^ = 0.31, 90% CI [0.18, 0.24]) and a smaller significant treatment-by-strain interaction effect for hypothermia (F21,232 = 2.81, p < 0.001, ωp^2^ = 0.11, 90% CI [0.00, 0.10]). A small but significant sex-by-treatment interaction was observed for the hypothermia trait (F1,232 = 6.81, p < 0.01, ωp^2^ = 0.02, 90% CI [0.00, 0.06]).

### Significant strain effects account for variation in initial responses to acute THC treatment

To demonstrate and visualize the direct effect of acute THC treatment in each BXD strain, we calculated the difference between day 1 and baseline day 0 as the initial THC response for each trait (**Figure 1B**). Based on the presence or absence of treatment-by-sex interaction effects, male and female initial THC responses were combined by strain for the mobility and analgesia traits and separated by sex for the hypothermia trait. Initial responses to THC measured in 20 BXD strains are summarized in **Figure 2**. The overall initial response distribution for each trait within males and females of each strain can be found in **Supplemental Figure 1**.

**Figure 2.**
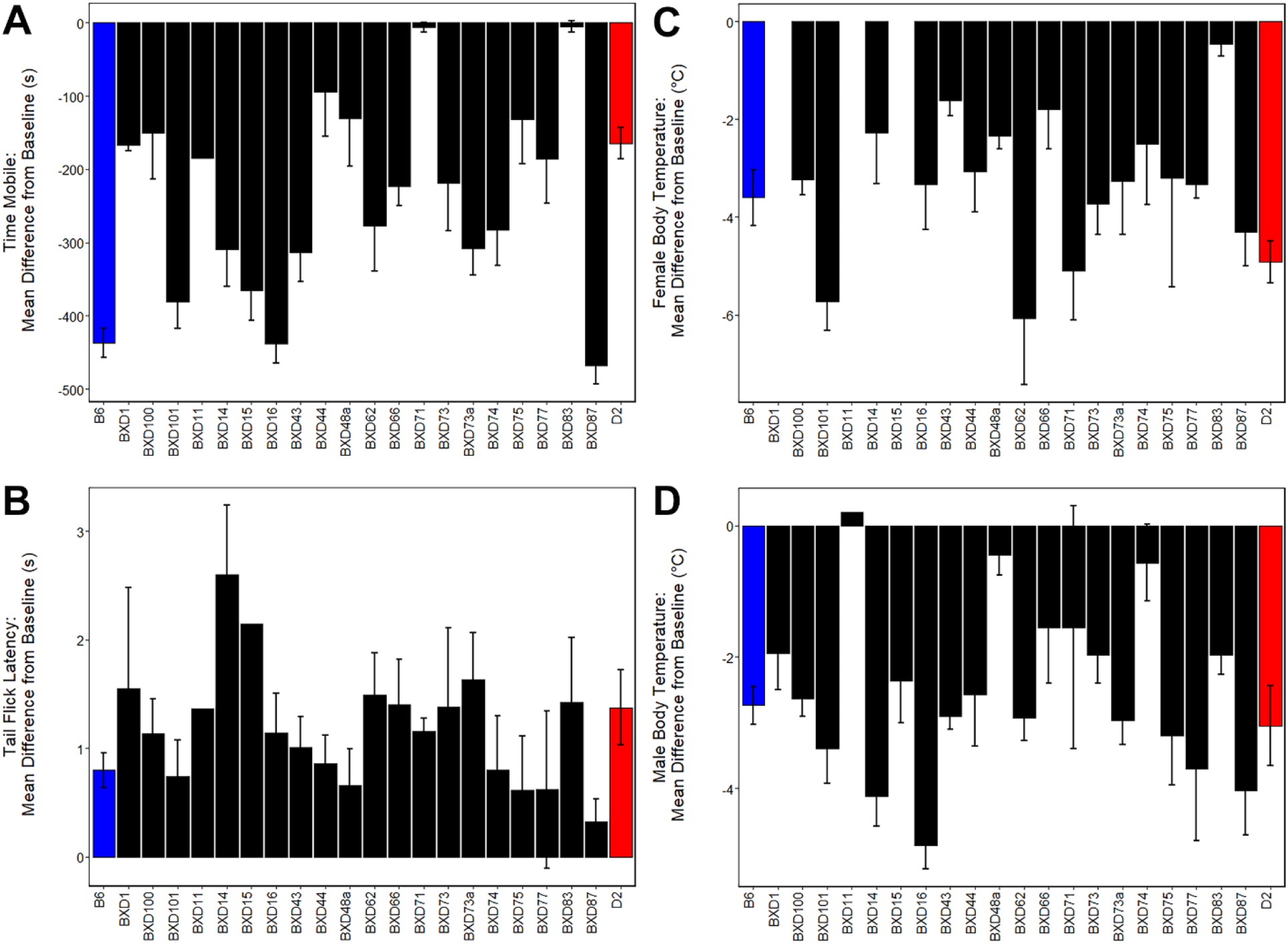
Variation in Initial Response to THC Among BXD and Parental Strains. A significant effect of THC treatment relative to baseline (*p* < 0.001) was observed for all traits. Initial response to THC is shown as the difference between baseline and initial THC treatment. Strains are shown on the x-axis and initial response (difference between baseline response on day 0 and response to THC on day 1) is shown on the y-axis. Negative values indicate a decrease in response on day 1 compared to day 0 and positive values indicate an increase in response. Parental strains are indicated in blue (B6) and red (D2) and BXD strains are shown in black. (A) There is a significant effect of strain (*p* < 0.001) on initial response to THC for time spent mobile in the open field 30 min post-injection. Males and female responses were combined as there were no significant interaction effects involving sex. Responses to THC varied ~90-fold among strains for the mobility trait. (B) The antinociceptive effects of THC were quantified using tail withdrawal latency in response to a thermal stimulus at 60 minutes post-injection. Males and female responses were combined as there were no significant interaction effects involving sex. Response to THC varied nearly 8-fold across strains, however, no significant effects of strain were observed for the analgesia trait. (C,D) There is a significant main effect of strain (*p* < 0.05) on initial hypothermic response 60 min post-injection of THC for both females (C) and males (D). Hypothermic response to THC in females varies 13-fold across strains compared to 24-fold variation in response in males.

For mobility, strain had a large and significant effect on initial response to THC (F21,121 = 9.01, p < 0.001, ω^2^ = 0.54, 90% CI [0.39, 0.59]). Average mobility across BXD strains was reduced by 238.40 ± 28.08 sec (all values reported as mean ± standard error, **Figure 2A**). Strains varied an astonishing 90-fold in their initial response to the immobilizing effects of THC. Similar to the sensitive B6 parental strain (–437.30 ± 19.49 sec), BXD16 (–439.70 ± 26.03 sec) and BXD87 (–468.13 ± 24.69 sec) demonstrated large reductions in mobility after acute THC. In contrast, BXD71 (–6.47 ± 6.47 sec) and BXD83 (–5.06 ± 7.50 sec) demonstrated lower sensitivity to THC, even when compared to the less sensitive D2 parental strain (–164.48 ± 21.14 sec).

Acute THC resulted in an average 1.20 ± 0.11 sec increase in tail withdrawal latency across all strains measured (**Figure 2B**). Initial analgesic/antinociceptive response to THC varied nearly 8-fold across strains with an increase in tail withdrawal latency from 2.60 sec in the most sensitive strain (BXD14) to only 0.33 sec in the least sensitive strain (BXD87). However, strain did not have a significant effect on initial analgesic response to THC (F21,129 = 1.25, p = 0.22, ω^2^ = 0.03, 90% CI [0.00, 0.00]).

Strain had a moderate and significant effect on initial hypothermic response to THC in females (F18,46 = 2.68, p < 0.01, ω^2^ = 0.32, 90% CI [0.00, 0.32]). In males the effect was also significant, but smaller (F21,66 = 1.78, p < 0.05, ω^2^ = 0.16, 90% CI [0.00, 0.03]). Average body temperature across BXDs was reduced by 3.36 ± 0.33 °C in female and by 2.42 ± 0.27 °C in males. BXD62 (−6.07 ± 1.35 °C) and BXD101 (−5.73 ± 0.60 °C) females and BXD16 (−4.86 ± 0.35 °C) males demonstrated greater relative sensitivity to the hypothermic effects of THC (**Figure 2C**). In contrast, BXD83 (−0.45 ± 0.25 °C) females and BXD11 (0.20 °C), BXD48a (−0.45 ± 0.30 °C), and BXD74 (−0.57 ± 0.58 °C) males demonstrated reduced sensitivity to the hypothermic effects of THC (**Figure 2D**). BXD females and males varied in initial response to the hypothermic effects of THC by 13-fold and 24-fold, respectively, and the range of responses in the BXDs exceeded that of the parental B6 and D2 strains.

### Variation in initial responses to THC are heritable among BXDs

Previously, we identified putative heritable strain differences in acute response to a single THC injection (10 mg/kg) in the parents of the BXD family—B6 and D2 (Parks et al. 2020). Here we test the hypothesis that polymorphic variants between these strains segregate among the BXD progeny and cause variation in THC initial responses. If responses to THC are variable and heritable among BXDs, then QTL mapping in the BXD population has the potential to identify causal variants in effectors of CB1R signaling.

To address this hypothesis, two different methods (*h*^2^ and ICC) based on the variance components of a one-way ANOVA were used to estimate heritability (**Table 2**). In general, THC response traits demonstrated high heritability. Tail withdrawal latency (a measure of analgesia) was an exception. Mobility and hypothermia traits demonstrated higher heritability, which indicated a strong contribution of genetic variants to phenotypic variation. Mobility and hypothermia traits also demonstrated sex differences with regards to heritability. Mobility had higher heritability in males whereas hypothermia had higher heritability in females.

### Initial response to the effects of THC on mobility and hypothermia in males are correlated and co-regulated by a locus on Chr 11

Examination of trait correlation structure can provide evidence of underlying co-regulation of trait expression due to biological factors. Based on the presence or absence of treatment-by-sex interaction effects, male and female initial THC responses were combined by strain for the mobility and analgesia traits and separated by sex for the hypothermia trait resulting in a total of four traits for correlation analysis (**Figure 3A**). We observed a significant (*p* < 0.01) and positive correlation between time mobile (males and females combined) and hypothermia in males, but not in females, following the first exposure to THC (**Figure 3A,D**). As expected, based on the presence of a strong treatment-by-sex interaction effect of THC, hypothermic responses to THC in males and females were modestly and positively correlated, but these correlations were not significant. Modest and positive (but not significant) correlations were also observed between the hypothermic response in females and time mobile following THC treatment. In contrast, THC-induced analgesia was uncorrelated with both the mobility trait, and female and male hypothermia traits.

**Figure 3.**
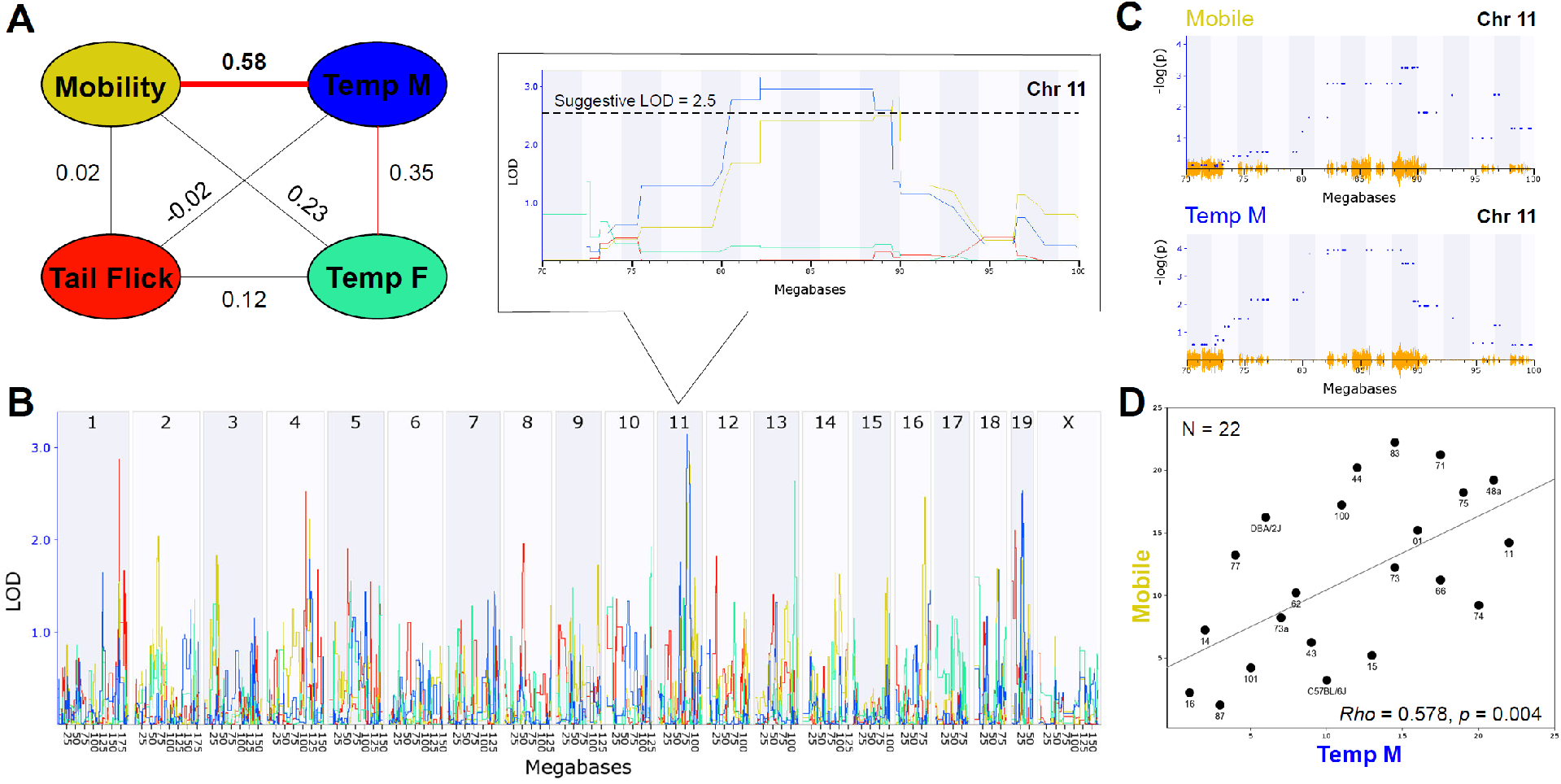
THC Initial Response Trait Correlations and Genetic Co-Regulation. (**A**) Trait correlation network constructed after calculating Pearson’s correlation on ranked trait values to generate Spearman’s rank order correlation coefficient (Rho or *P*). The hypothermia trait was split by sex due to a condition-by-sex interaction effect on body temperature change in response to THC. Changes in time mobile in the open field and body temperature in males following a single THC (10 mg/kg) exposure are significantly (p < 0.01) correlated. Weak (not significant) correlations are also observed between mobility and hypothermia in females following a single THC exposure and between changes in male and female hypothermia in response to THC. (**B**) Genetic regulation of THC initial response traits is observed, albeit at a suggestive level with no traits passing the threshold for genome-wide significance (*p* < 0.05). Interval maps for each trait shown as a different line color corresponding to the correlation network in (**A**). The strength of association (LOD) on the y-axis is plotted for each trait across the genome (megabase position on each chromosome or Chr) on the x-axis. Change in mobility and male body temperature are regulated by the same suggestive locus on Chr 11 (zoomed region in boxed area). (**C**) Use of an alternative linear mixed-model QTL mapping method (GEMMA, see methods) resulted in replication of genetic co-regulation of initial motor and hypothermia THC response traits from the same Chr 11 interval. Strength of association or −log(P) values (y-axis) shown for each marker (blue dots) within the Chr 11 QTL interval for mobility and temperature (males) initial THC response traits. Mapping with linear mixed-models can account for population family structure or kinship within the BXD population that is not addressed using traditional interval mapping. SNP density plotted in orange on the x-axis. (**D**) Scatterplot describing the relationship between average change in time mobile in the open field in response to THC (X-axis) and average change in male body temperature in response to THC (Y-axis) for each BXD strain based on Spearman’s rank order correlation.

Heritability estimates were high for several initial THC response traits (*i.e*. change in time mobile in the open field in response to THC; **Table 1**), indicating strong regulation of trait expression by genetic variants. To explore the potential for future genetic mapping studies of these traits in a larger BXD panel, we performed QTL scans for all traits. As expected, no genome-wide significant QTLs (LOD > ~3.5) were detected for any trait (**Figure 3B**) using traditional interval HK mapping methods. However, suggestive QTLs (LOD > 2.5) were detected for all THC initial response traits (**Table 2**). Moreover, the highly correlated mobility and male hypothermia traits were co-regulated by a QTL on Chr 11 with a 1.5 LOD drop confidence interval between 80 and 90 Mb (**Figure 3B**).

**Table 1.**
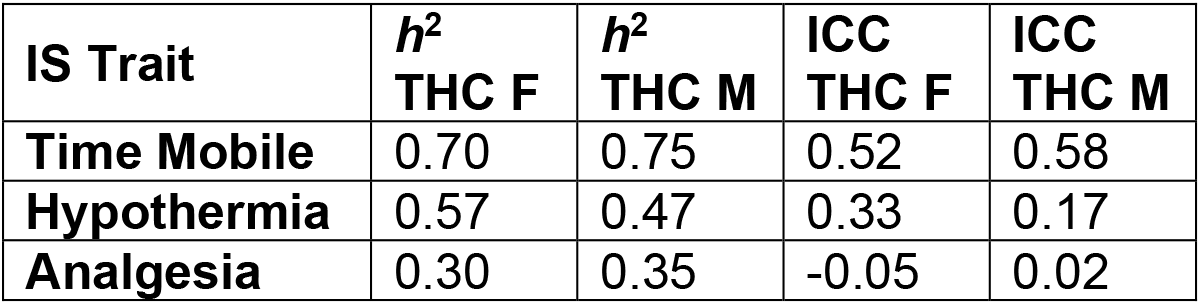
Estimates of Heritability Among BXD and Parental Strains. IS = initial sensitivity; *h*^2^ = narrow sense heritability; ICC = intraclass coefficient estimate of heritability; F = female; M = male.

| IS Trait | $h^2$<br>THC F | $h^2$<br>THC M | ICC<br>THC F | ICC<br>THC M |
| --- | --- | --- | --- | --- |
| Time Mobile | 0.70 | 0.75 | 0.52 | 0.58 |
| Hypothermia | 0.57 | 0.47 | 0.33 | 0.17 |
| Analgesia | 0.30 | 0.35 | -0.05 | 0.02 |

**Table 2.**
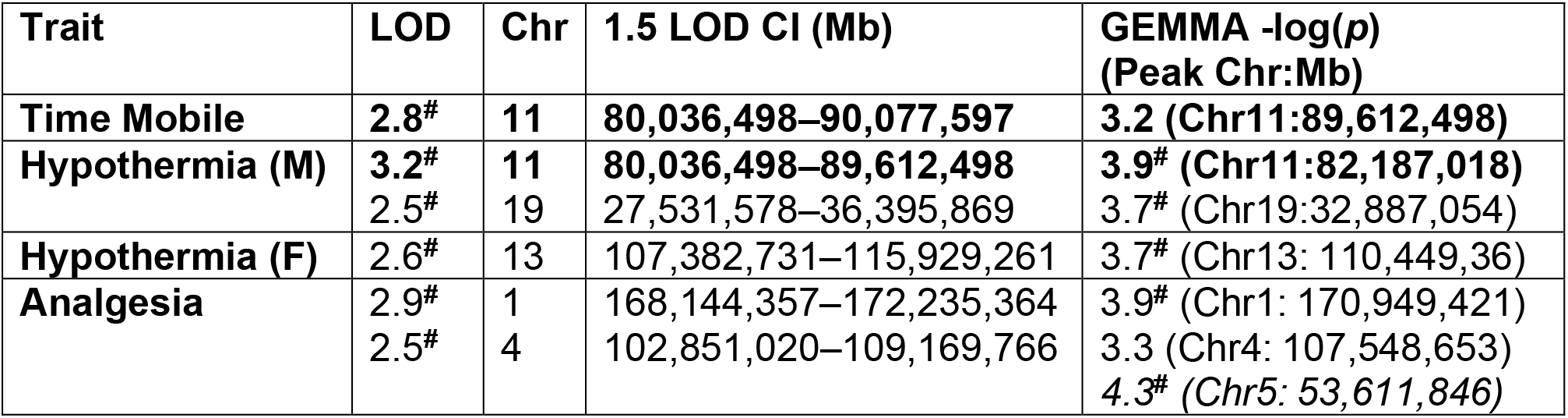
Summary of Suggestive QTLs for THC Initial Response Traits. LOD = logarithm of the odds score based on traditional simple regression; Chr = chromosome; CI = confidence interval; # = meets suggestive genome-wide threshold; F = female; M = male. Bold text indicates overlapping QTL shared by mobility and hypothermia (male) traits and italicized text indicates QTL detected only using GEMMA.

Shared genetic co-regulation of mobility and male hypothermia traits on Chr 11 was also replicated using a different QTL mapping method (GEMMA, **see Methods**) that leverages linear mixed-models and accounts for population family structure (**Figure 3C**). Suggestive QTLs for initial response to the hypothermic (Chr 19 in males and Chr 13 in females) and analgesic (Chrs 1 and 4) effects of THC also replicated across both QTL mapping methods (**Table 2**). However, the mobility (Chr 11) and analgesia QTL (Chr4) did not meet the suggestive threshold using GEMMA. Additional suggestive QTLs for analgesia (Chr 5) were detected by GEMMA only (**Table 2**).

### Effectors of endocannabinoid signaling identified as positional candidates for THC initial response QTLs

As a first step towards identification of variants and genes modulating initial responses to THC, we prioritized several suggestive QTLs for exploration of putative candidate genes. From **Table 2**, we selected the Chr 11 locus modulating both mobility and male hypothermia in response to THC and loci on Chrs 19 (male hypothermia), 13 (female hypothermia), Chr 1 (analgesia), and Chr 4 (analgesia). All QTLs were replicated using both HK and GEMMA mapping methods. Following a standard workflow (**see Methods**) we identified a number of genes near or overlapping SNPs, InDels, or SVs, some of which are predicted to have an impact on transcript or protein integrity (*e.g*., missense, stop gain or loss, splice region variant, or frameshift) (**Supplemental Table 1**). Based on evidence in support of genetic regulation of gene expression by cis eQTLs in naive BXD brain tissue (**Supplemental Table 2**) and literature associations with relevant cannabinoid terms detected by RatsPub (**see Methods**), we nominated several positional candidates for each prioritized QTL. Top positional candidates include *Ndufs2* (Chr 1 Analgesia QTL); *Scp2* (Chr 4 Analgesia QTL); *Rps6kb1*/p70s6K (Chr 11 Mobility and male Hypothermia QTL); *Pde4d* (Chr 13 female Hypothermia QTL); and *Pten* (Chr 19 male Hypothermia QTL) (**Table 3** and **Supplemental Table 3**). These genes are likely to play a role in modulating initial response to THC based on location within modulatory QTL, putative functional sequence variants between B and D haplotypes that segregate among BXD progeny, evidence of genetic control of expression in brain tissue, and previously reported involvement in endocannabinoid/cannabinoid receptor signaling pathways. Below we summarize the evidence in support of these top positional candidates.

**Table 3.**
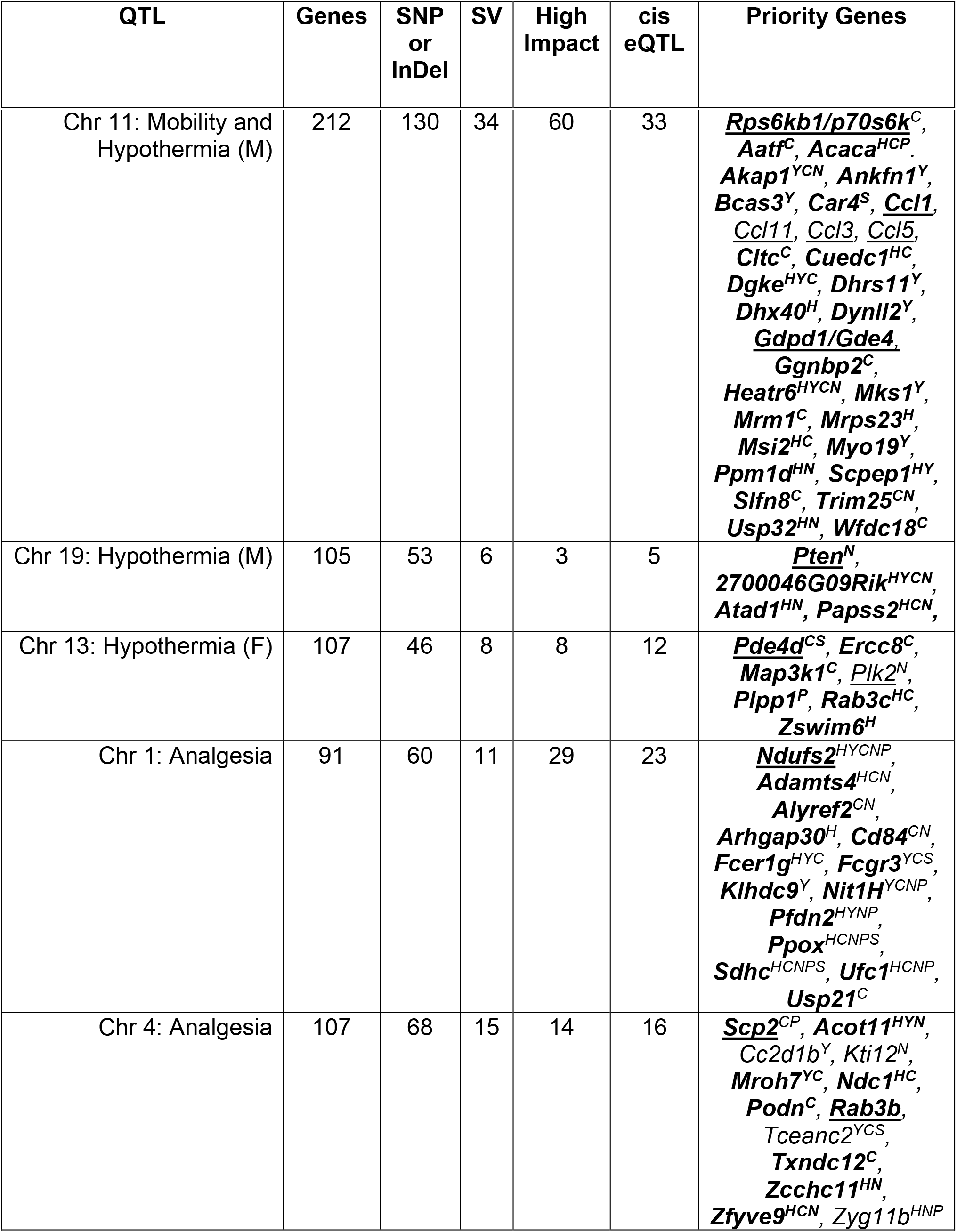
Summary of Genes with Variants and High Impact Variants Within QTL CIs. Number of genes or noncoding RNA (ncRNA) within each interval (*Genes* column), number of genes or ncRNA near or overlapping a SNP or InDel (SNP or *InDel* column), number of genes or ncRNA overlapping a SV (*SV* column), and number of genes or ncRNA containing a high impact SNP or InDel (*High Impact* column) are shown for each THC initial response QTL. Number of genes within each QTL CI whose expression is regulated by a cis eQTL are shown in the *cis eQTL* column. High priority candidate QTL genes (QTGs) associated through literature searches with endocannabinoid/cannabinoid signaling whose expression (underline), regulated by a cis eQTL in at least one brain region (Y = hypothalamus, C = neocortex, S = striatum, N = nucleus accumbens, P = prefrontal cortex) and/or overlap a SV or high impact SNP or InDel (bold) are shown in the *Priority Genes* column.

#### NADH dehydrogenase [ubiquinone] iron-sulfur protein 2 (Ndufs2) is a positional candidate for modulation of the analgesic response to THC

*Ndufs2* encodes a core subunit of the mitochondrial Complex 1. The gene also contains both missense and putative splice region variants and demonstrates evidence of genetic regulation of expression in the form of cis eQTLs detected in naive BXD hippocampus, hypothalamus, neocortex, nucleus accumbens, and prefrontal cortex. Moreover, *Ndufs2* has been implicated in mediating response to cannabinoids. Activation of mitochondrial CB1Rs resulted in a decrease in PKA-dependent phosphorylation of oxidative phosphorylation proteins, including NDUFS2, and a concomitant decrease in brain mitochondrial function that was associated with cannabinoid-induced synaptic depression and amnesia (Hebert-Chatelain et al. 2016).

#### Sterol carrier protein 2 (Scp2) is a positional candidate for modulation of the analgesic response to THC

The SCP2 protein has been proposed to regulate brain endocannabinoid levels and, thus, endocannabinoid system function (Liedhegner et al. 2014; Martin et al. 2019). The *Scp2* gene contains a putative splice region variant and demonstrates genetic modulation of expression (cis eQTL) in naive BXD neocortex and prefrontal cortex.

#### Ribosomal Protein S6 kinase (Rps6kb1) is a positional candidate for modulation of the effects of THC on both mobility and hypothermia in males

The gene *Rps6kb1* encodes the 70-kDa ribosome protein S6 kinase (p70S6K) and contains a predicted splice region variant. Expression of *Rps6kb1* is modulated by a cis eQTL in neocortex. Activation of the G-protein coupled CB1R by THC triggers cascades of intracellular signaling, including activation of phosphoinositide-3 kinase (PI3K)/Akt/glycogen synthase kinase 3 (GSK-3) and subsequent activation of the serine/threonine kinase mammalian target of rapamycin (mTOR). P70S6K is a downstream target of mTOR whose activation is associated with protein synthesis and regulation of different cellular states (*e.g*., survival, growth, or autophagy). The CB1R/mTOR/p70S6K signaling pathway in hippocampal GABAergic interneurons was found to mediate THC-induced long-term memory deficits (Puighermanal et al. 2009; Puighermanal et al. 2013).

#### Phosphodiesterase 4D, cAMP specific (Pde4d) is a positional candidate for modulation of the hypothermic response to THC in females

Phosphodiesterase 4 hydrolyzes cAMP and plays a role in mediating cAMP signaling pathways and neuroadaptations favoring drug reinforcement, tolerance and dependence (Nestler 2016; Cherry and Davis 1999; Perez-Torres et al. 2000; Muschamp and Carlezon 2013). Phosphodiesterase inhibitors have shown some efficacy in reducing drug seeking behavior or intake of psychostimulants, alcohol, and opioids in preclinical models (reviewed in (Olsen and Liu 2016)). There are four isoforms of Phosphodiesterase 4, including the positional candidate PDE4D, which expresses 11 alternative splice variants and exhibits widespread expression in brain (Olsen and Liu 2016). Expression of *Pde4d* is modulated by cis eQTLs in naïve BXD neocortex and striatum and the gene locus contains a missense variant as well as a putative splice region variant.

#### Phosphate and tensin homolog (Pten) is a positional candidate for the modulation of the hypothermic response to THC in males

Highly expressed in brain, *Pten* encodes a dual protein and lipid phosphatase enzyme that has been proposed to interact with neurotransmitter receptors, including NMDA receptors (Ning et al. 2004) and serotonin 5-HT2C receptors (Ji et al. 2006), and regulate their activity. Disruption of the interaction between PTEN and serotonin 5-HT2C receptors in the ventral tegmental area (VTA) has been shown to inhibit the firing rate of dopaminergic VTA neurons projecting to the nucleus accumbens (Ji et al. 2006). Moreover, disruption of this interaction also blocked the ability of THC to enhance the firing rate of dopaminergic VTA neurons and subsequent THC-induced conditioned place preference (Ji et al. 2006) suggesting that interactions between PTEN and serotonin receptors may mediate some of the behavioral effects of cannabinoids and other drugs of abuse (Maillet et al. 2008). The expression of *Pten* in naive BXD nucleus accumbens is modulated by a cis eQTL and there are 59 variants located within or near the gene locus, although the impact of these variants on gene function or expression is not clear.

### Positional candidates identified as putative modulators of initial responses to THC have been previously associated with drugs of abuse

For each high priority positional candidate or QTL gene (QTG) we explored broader associations with over 300 addiction and psychiatric disease key words using a newly developed resource: *Relationship with Addiction Through Searches of PubMed* (*RatsPub*, **see Methods**). This search finds sentences in PubMed that contain both the gene of interest and keywords relevant to drugs of abuse, addiction, and psychiatric disease. Our overall goal was to place provisional endocannabinoid signaling QTGs identified in our study into a broader biological context. Given the broad modulatory effects of the endocannabinoid system on brain function, we hypothesized that THC initial response QTGs would also be associated with diverse signaling in response to other drugs of abuse and psychiatric disorders.

*Pten* (Chr 19 male Hypothermia QTG), *Rps6kb1*/p70S6K (Chr 11 Mobility and male Hypothermia QTG), and *Pde4d* (Chr 13 female Hypothermia QTG) were the most well connected QTGs in terms of associations with drugs of abuse and psychiatric disease (**Supplemental Figure 2**). All three of these QTGs are associated with terms related to stress, anxiety, depression, and schizophrenia. *Rps6kb1*/p70S6K and *Pten* were associated with terms related to alcohol, amphetamine, cocaine, nicotine, opioid, phychedelics, and sensitization. *Pde4d* was associated with terms related to nicotine and psychedelics. *Pten, Rps6kb1*/p70S6K, and *Pde4d* were also associated with neuroplasticity, neurotransmission, signaling and transcription terms in support of their known roles in these biological processes. Moreover, these three QTGs have also been implicated in brain regions relevant to drug abuse and psychiatric disease as evidenced by their literature co-citation with relevant terms (*e.g*., accumbens, amygdala, cortex, habenula, hippocampus, or striatum).

Although less well-connected, the QTGs *Ndufs2* (Chr 1 Analgesia QTL) and *Scp2* (Chr 4 Analgesia QTL) were also associated with neuroplasticity, signaling, transcription, stress, and hippocampus terms (**Supplemental Figure 2**). In addition, *Ndufs2* was associated with schizophrenia terms and *Scp2* was associated with alcohol-related terms.

### Initial responses to THC are correlated with initial responses to other drugs of abuse and cocaine self-administration

Recent studies quantified initial response to several drugs of abuse (*i.e*. cocaine, alcohol, morphine) or intravenous cocaine self-administration among a large set of BXDs (Philip et al. 2010; Dickson et al. 2016). BXD traits associated with these studies are available in GN and are of sufficient strain depth (see **Methods**) to facilitate a moderately well-powered and exploratory correlation analysis with our THC initial response traits. We hypothesized that strong correlations between initial responses to THC and initial response or self-administration of other drugs of abuse among BXD strains was indicative of shared genetic and/or biological co-regulation.

From our background list of 302 BXD legacy traits, we found 43 significant (*p* < 0.05) correlations (**Supplemental Table 3**) between 40 drug response or self-administration traits and one or more of our THC initial response traits. Several interesting correlation patterns were evident between initial responses to THC and BXD legacy cocaine, ethanol, and morphine response traits (**Table 4**). One of the most striking patterns was an enrichment of significant and positive correlations between THC-induced analgesia and initial locomotor responses to morphine (50 mg/kg i.p.) from 90 to 150 min post-injection. Other patterns emerged as well: (1) significant and positive correlations between THC-induced hypothermia (females) and naloxone-induced (30 mg/kg i.p.) withdrawal from morphine (50 mg/kg i.p.), (2) significant and positive correlations between cocaine self-administration and THC-induced immobility, (3) sex-specific and significant correlations between THC-induced hypothermia and preference for cocaine, and (4) significant correlations between both THC-induced immobility and hypothermia (females) and initial motor response and tolerance to alcohol/ethanol.

**Table 4.**
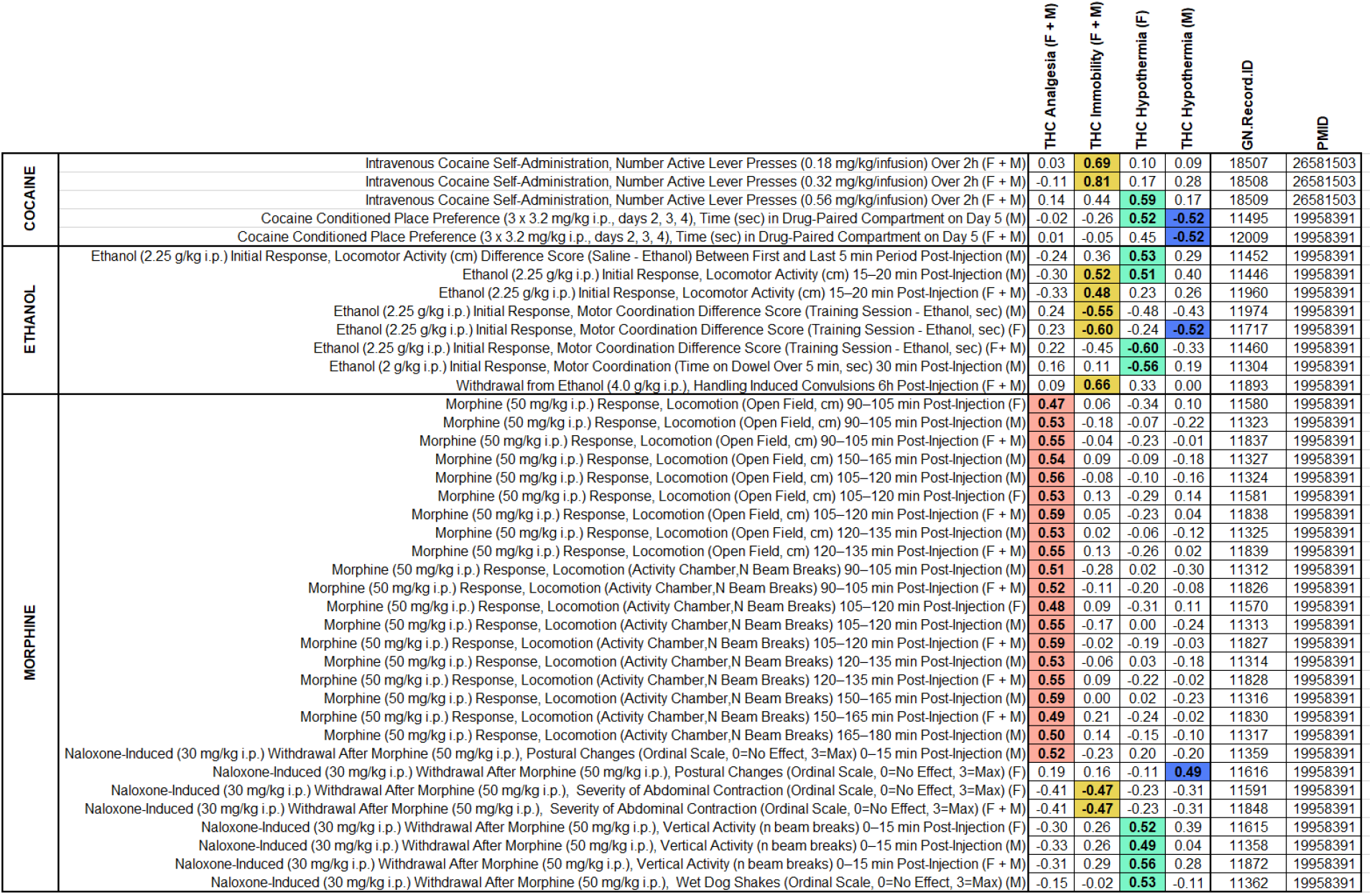
Summary of Significant (p < 0.05) Correlations Between Initial Response to THC and BXD Legacy Drug Response Traits.

The role of sex on trait covariation was not explicitly quantified in this exploratory analysis. However, strong correlations were detected between each THC trait and BXD legacy trait measurements both when sex was combined or considered separately. Finally, correlation patterns did not appear to be driven by genetic co-regulation of cannabinoid and drug response traits from shared loci (*i.e*. peak QTLs for correlated traits were non-overlapping). Instead, these correlation patterns indicate shared biological mechanisms and more complex genetic modulation driving response to THC and other drugs of abuse.

## DISCUSSION

In this study we demonstrate that strain differences in initial responses to THC between B6, D2, and their BXD progeny are heritable (**Figure 2** and **Table 1**). Thus, genetic mapping of THC response traits in the BXD family will lead to the identification of genomic loci, including variants and genes, that modulate response to cannabinoids. Candidate QTGs will almost certainly be direct or indirect effectors of endocannabinoid signaling given that the THC response traits measured in our study are entirely downstream of CB1R signaling. Following this line of reasoning, we identified multiple QTLs and positional candidate endocannabinoid signaling QTGs (*Pten, Rps6kb1*/p70S6K, *Pde4d, Ndufs2, Scp2*) that may control variation in initial response to THC (**Tables 2, 3**). Many of these candidates mediate responses to other drugs of abuse (**Supplemental Figure 2**). Moreover, we found significant correlations between initial responses to THC and behavioral responses to cocaine, alcohol, and morphine (**Table 4**). Taken together, we provide strong evidence that gene variants in endocannabinoid signaling pathway genes are responsible for individual variation in initial THC responses and are likely to cause variation in the behavioral responses to other drugs of abuse.

Our genetic screen for natural variation in traits modulated by effectors of endocannabinoid signaling was designed to identify specific genes and pathways involved in the response to THC. Targeted genetic deletion has revealed insight into the integral role of CB1R in this response. However, little is known about the role of natural variation in response to cannabinoids. A handful of genes (*e.g. CADM2, NCAM1, NRG1, CSMD1*, and *CHRNA2*) have been associated with cannabis use or dependence in human genome-wide association studies (Demontis et al. 2019; Han et al. 2012; Pasman et al. 2018; Sherva et al. 2016; Stringer et al. 2016). The underlying mechanisms whereby these genes mediate use or dependence to cannabinoids or interact with the endogenous cannabinoid system has yet to be elucidated, with one notable exception. Heterozygous *Nrg1* (Neuregulin) mutant mice, in which one copy of the gene contains a deletion of the transmembrane region, are more sensitive to the initial locomotor suppressant and behavioral effects of THC (Boucher et al. 2007) and develop more rapid tolerance to the motor suppressant effects (Boucher et al. 2011) relative to control mice. In naïve heterozygous *Nrg1* mutant mice relative to controls, these differences in cannabinoid responses were preceded by modest increases of CB1R levels in substantia nigra and significant decreases in both thalamic NMDA receptor levels and striatal dopamine D2 receptor levels (Newell, Karl, and Huang 2013). The association of *NRG1* variants with cannabis dependence in humans and independent preclinical evidence for a role of *Nrg1* in initial response to cannabinoids is striking. Identification of additional genes that modulate endocannabinoid signaling and cannabinoid response is non-human genetic populations is clearly warranted and could help clarify associations determined in humans.

To date, no human studies have addressed the impact of genetic variation on acute behavioral responses to cannabis or derived cannabinoids. Greater understanding of the genes and pathways mediating responses to cannabinoids is important for at least two reasons. First, identification of variants in endocannabinoid signaling pathways will help us better understand biological responses to cannabinoids, including risk of cannabinoid dependence, and provides the opportunity to decouple unwanted side effects with therapeutic properties of cannabinoid drugs (*e.g*., motor impairment versus analgesic effects of THC). Second, endocannabinoid pathways represent common points of convergence that are highly relevant for understanding the molecular response to many drugs of abuse and molecular mechanisms underlying addiction processes. To this end, we have identified strong correlations between initial responses to THC and initial response or self-administration of cocaine, morphine, and/or alcohol among BXD strains that could be indicative of shared genetic and biological co-regulation.

The endocannabinoid system (*e.g*., lipid ligands, biosynthetic and catabolic enzymes, G-protein coupled receptors and effectors of signaling) plays an important role in modulating synaptic plasticity, behavior, and reward circuitry. Cannabis, cannabinoids and non-cannabinoid drugs of abuse have been shown to disrupt endocannabinoid system function and signaling, and these changes may contribute to addiction processes (Parsons and Hurd 2015). For example, exposure to alcohol and opioids has been shown to increase the level of endocannabinoid system ligands (AEA or 2-AG) in rodent brain reward regions (Alvarez-Jaimes, Polis, and Parsons 2009; Alvarez-Jaimes, Stouffer, and Parsons 2009; Caille et al. 2007; Ceccarini et al. 2013). Moreover, activation of CB1Rs in some rodent models has been shown to enhance the rewarding effects of alcohol and opioids through both dopamine dependent and independent pathways (Ellgren, Spano, and Hurd 2007; Serrano and Parsons 2011; Panagis, Mackey, and Vlachou 2014). Both cocaine (Grueter et al. 2006; Liu, Pu, and Poo 2005; Pan, Hillard, and Liu 2008; Fourgeaud et al. 2004) and alcohol (Clarke and Adermark 2010; Adermark et al. 2011) exposure have been shown to alter CB1R-mediated plasticity in rodent brain reward regions (e.g., striatum, ventral tegmental area, nucleus accumbens, and/or bed nucleus of the stria terminalis). CB1 R-mediated plasticity and signaling is also important for mediating response to opioids (Guegan et al. 2016; Zhao et al. 2017). Thus, the endocannabinoid system converges at many points with other neurotransmitter systems to mediate drug response and alter behavior following drug exposure.

We hypothesized that correlations between initial response to THC and responses to alcohol, cocaine, or opioids in the BXD population resulted from the convergence of genes and pathways that mediate both effects. Indeed, we observed striking correlations between THC and opioid responses (enhanced sensitivity to the analgesic effects of THC and enhanced sensitivity to the motor stimulant effects of morphine), THC and cocaine responses (enhanced sensitivity to the motor suppressant effects of THC and higher active lever presses for cocaine), THC and ethanol responses (enhanced sensitivity to the motor suppressant and hypothermic effects of THC and enhanced sensitivity to the motor stimulant effects of ethanol), and THC and withdrawal from alcohol and opioids (enhanced sensitivity to the motor suppressant effects of THC and higher levels of withdrawal from alcohol in contrast to lower levels of precipitated withdrawal from opioids). The results of these exploratory correlational analyses hint at underlying shared genetic and biological regulation. Of interest, no single shared locus could account for the covariance in drug response traits, indicating complex regulation by multiple loci, genes, and variants. Importantly, our exploratory analysis identified individuals of the BXD population with extreme responses to multiple drugs of abuse. These individuals can be leveraged to directly test predictions about drug response and to identify mediators of behavioral responses to both cannabinoid and non-cannabinoid drugs using both genetic (QTL mapping) and pharmacological (through targeting of specific pathways) methods.

Our study marks the first and largest attempt to quantify genetic factors mediating initial responses to THC in males and females of a genetic population. While we have made an important contribution to the field, there are some limitations of our study that will need to be addressed by future experiments. First, THC traits were profiled in a subset of 20 BXD strains which limits our ability to detect QTLs of small effect size. All QTLs identified in our study are suggestive after accounting for the effects of genome-wide testing. It is likely that these loci will be replicated in an independent or larger cohort of BXD strains. We emphasize that this replication will be required to confirm QTLs and QTGs prioritized by our study. Other aspects of our study design limit translatability somewhat and should be addressed by future studies. These include the use of a single major component of cannabis (THC), use of a single dose (10 mg/kg), and quantification of acute physiological responses to THC (as opposed to chronic exposure, dependence and/or withdrawal). Finally, we were agnostic to genetic variation in cannabinoid metabolic pathways in this study.

Drug metabolism is an important biological process that impacts response to drugs of abuse and susceptibility to addiction. Based on trait correlation structure and genetic mapping, it is unlikely that strain differences in THC metabolism contributed to trait variation in all three of our initial response traits (*e.g*., mobility, hypothermia, and analgesia). It will be important in the future to establish whether sex or strain differences in THC metabolism contribute to variation in response and whether there are functional variants that impact THC metabolism in rodents. Several functional human variants (alleles) have been identified for cannabinoid metabolism genes (e.g. CYP3A4 and CYP2C9, reviewed in (Hryhorowicz et al. 2018)). The impact of these variants on response to cannabinoids and addiction is not clear, although a small study reported a trend towards increased sensitivity to THC in individuals homozygous for the CYP2C9*3 allele associated with slower clearance of THC (Sachse-Seeboth et al. 2009).

Ultimately, our study demonstrates the feasibility of genetic mapping in the BXDs to identify underlying biological and genetic factors contributing to response to THC and other drugs of abuse. We have taken important first steps towards the identification of loci and genes that modulate initial responses to THC. We have also provided evidence of covariation among drug response traits, although the underlying causal factors remain elusive. Finally, we have identified BXD individuals with extreme responses to acute THC exposure. Identification of BXD individuals with greater initial sensitivity to THC will facilitate more detailed genetic analysis of intake, reward, withdrawal, tolerance, and the behavioral impact of cannabinoids on motivation, cognition, and health. The genetic architecture of behavioral and pharmacological responses to cannabis and cannabinoids is difficult to reconstruct using currently available human data sets. A better understanding of trait correlation structure and regulation by underlying genetic variation in diverse rodent models is expected to help bridge these gaps and direct the focus of future human studies.

## METHODS

### Experimental subjects

The BXD family (**Figure 1A**) is maintained in a large breeding colony at the University of Tennessee Health Science Center. Male and female mice from 20 strains currently breeding in the colony were made available upon weaning. Each strain was composed of mice from one or two litters. Age ranges on the first day of testing were from 57 to 154 days-of-age. Mice were assigned to testing cohorts consisting of a maximum of 32 mice. Due to the number of strains and availability in the breeding colony, complete counterbalancing of strain, sex and condition across cohorts was not practical. At least one week prior to testing, animals were housed individually and handled daily. Handling consisted of lifting each individual in either cupped hands (avoiding any lifting by the tail) or a plastic lid from a pipette tip box. Food and water were provided *ad libitum* and animals were maintained on a 12 h:12 h light:dark cycle. All testing was performed during the light cycle from 0700 to 1600 hours. All procedures were approved by the University of Tennessee Health Science Center Institutional Animal Care and Use Committee.

### THC and vehicle formulation

THC was formulated in an ethanol:cremophor:saline (5:5:90) vehicle followed by filter sterilization. The resulting formulation was stored in the dark at 4°C in a septum sealed vial. Vehicle (VEH) was prepared in the same manner. THC and VEH were administered by intraperitoneal (*i.p*.) injection at a dose of 10 mg/kg such that a 30 g mouse received a 100 μl injection volume. Each formulation was used for the entire testing schedule of each cohort (see details below). Previously we demonstrated that each THC formulation contains 95.2% of the initial amount of THC when measured eight days after preparation (Parks et al. 2020).

### Testing schedule

Treatment and trait measurements are described in **Figure 1B,C.** Mice of each sex and strain were included (see **Table 5** for exact numbers). On the first day (Day −1), no injections were given. On the second day (Day 0, baseline), all mice received an injection of VEH (100 μl per 30 g). On Day 1, mice received an injection of 10 mg/kg THC. On each day, body temperature, tail withdrawal latency in response to a thermal stimulus, and activity (time mobile) in the open field (OF) were measured at multiple time points in the same animal. Rectal body temperature was measured using a ThermoWorks digital thermometer with a mouse rectal probe adaptor at time 0 and 60 min post-injection. Tail withdrawal latency was measured 60 min post-injection by gently restraining each mouse in a 50 mL conical tube. The tail was then submerged ~2 cm into a 52 °C water bath. The latency to remove the tail was recorded to the nearest second. In most cases, tail flick latency was measured by two independent experimenters, and the resulting scores were averaged. Time spent mobile or immobile (periods of no movement lasting for 3 or more secs) was measured 30 min post-injection in a 40 x 40 x 40 cm OF over a 10 min interval using video recording and ANY-maze (Stoelting) tracking software.

**Table 5.** Summary of Experimental Subjects.

| Strain | Sex | # |
| --- | --- | --- |
| B6 | M | 17 |
|  | F | 9 |
| D2 | M | 17 |
|  | F | 6 |
| BXD1 | M | 2 |
|  | F | 0 |
| BXD11 | M | 1 |
|  | F | 0 |
| BXD14 | M | 5 |
|  | F | 3 |
| BXD15 | M | 3 |
|  | F | 0 |
| BXD16 | M | 3 |
|  | F | 3 |
| BXD43 | M | 2 |
|  | F | 5 |
| BXD44 | M | 3 |
|  | F | 3 |
| BXD48a | M | 4 |
|  | F | 3 |
| BXD62 | M | 3 |
|  | F | 3 |
| BXD66 | M | 2 |
|  | F | 3 |
| BXD71 | M | 2 |
|  | F | 2 |
| BXD73 | M | 3 |
|  | F | 3 |
| BXD73a | M | 3 |
|  | F | 3 |
| BXD74 | M | 3 |
|  | F | 3 |
| BXD75 | M | 3 |
|  | F | 3 |
| BXD77 | M | 3 |
|  | F | 3 |
| BXD83 | M | 3 |
|  | F | 2 |
| BXD87 | M | 3 |
|  | F | 3 |
| BXD100 | M | 3 |
|  | F | 3 |
| BXD101 | M | 3 |
|  | F | 4 |

### Statistical analysis of THC treatment effects

Data from day 0 (VEH treatment) and day 1 (THC treatment) were used in a three-way between-subjects (treatment*strain*sex) analysis of variance (ANOVA). The purpose of this analysis was twofold: we wanted to determine if THC treatment had a large and significant effect on responses and quantify any interactions between treatment and sex that would justify additional sex-specific analyses. ANOVA was performed using base functions in R (*anova.lm* function). Effect sizes (partial omega-squared or ωp^2^) for each parameter in the ANOVA were calculated in R using the *omega_squared* function in the effectsize package. Response during habituation on day –1 was not analyzed and individual BXD data points were only removed for technical reasons (*e.g*. equipment failure during recording).

### Statistical analysis of initial responses to THC

Because THC treatment effects were significant and of moderate to large effect for all traits, initial responses to THC were calculated as the difference between day 1 and day 0 (baseline) for each trait (**Figure 1B**). Difference scores provide a more direct method to demonstrate initial THC responses and are more robust to the effects of batch (note that counterbalancing was not possible in the study design). Individual difference scores (initial response) data points for the parental B6 and D2 strains were excluded from further analysis if they were found to be two standard deviations above or below the mean trait values. The low number of replicates among BXD strains precluded outlier analysis (**see Table 5**). Male and female responses were combined for the analgesia and mobility initial sensitivity traits and separated for the hypothermia initial sensitivity trait based on detection of a significant sex-by-treatment interaction effect for the hypothermia in the three-way ANOVA to evaluate THC treatment effects. Strain averaged initial sensitivity traits for mobility and analgesia (Record IDs 21481 and 21483, respectively) and strain and sex averaged initial sensitivity for the hypothermia trait (Record IDs 21485 and 21487) have been uploaded to the GeneNetwork (GN, www.genenetwork.org) web service (RRID:SCR_002388) for the *Mouse* species, the *BXD Family* group, the *Traits and Cofactors* type, and the *BXD Published Phenotypes* dataset.

To quantify and visualize the effect of strain on initial response to THC for each trait (*y*), one-way between-subjects ANOVA in the form of *y* ~ strain was performed using base functions in R (*anova.lm* function). Effect sizes (omega-squared or ω^2^) for each ANOVA were calculated in R using the *omega_squared* function in the effectsize package.

### Heritability

Heritability is an estimate of the proportion of the trait variation that can be explained by genetic factors. Heritability was estimated for each sex using two different methods based on the variance components of a one-way ANOVA (y ~ Strain). First, narrow sense heritability (*h*^2^) was calculated by comparing the between-strain variance with the total variance (between- and within-strain or environmental variance (Hill and Mackay 2004)). This can be calculated from an ANOVA results table by dividing the between-strain sum of squares (SS strain) term by the sum of the between-strain and within-strain (total) SS terms (Lariviere and Mogil 2010). Second, the intraclass coefficient estimate of heritability (ICC) was estimated using the intraclass correlation coefficient (*ICCest* function in the ICC package in R) to determine how closely individuals within the same strain resemble each other.

### Genetic Mapping

Two different methods were applied to compute quantitative trait loci (QTL) probability given strain genotypes and initial responses to THC using GN (Mulligan et al. 2017). The first method applied was traditional simple regression (Haley-Knott or HK) and the second was genome-wide efficient mixed model association [GEMMA; (Zhou and Stephens 2012)] with the “leave one chromosome out” (LOCO) option [rationale for using both methods described in (Mulligan et al. 2018)]. Prior to performing genome scans, each trait distribution was checked for normality using the *probability plot* function in GN. All raw trait data were approximately normally distributed and used for QTL mapping. For both HK and GEMMA, a dense panel of 7,321 markers was used for mapping. QTL were considered noteworthy if: (1) a genome-wide suggestive level (*p* < 0.63, equivalent to a 63% probability of a false positive or one false positive per genome scan) was reached following permutation (1000 tests) using the HK method, and (2) the peak marker association [-log(p) > 3] detected using GEMMA overlapped the QTL mapped by HK.

GEMMA permutation was performed for each *BXD Published Phenotypes* dataset trait as described previously (Mulligan et al. 2018). For trait 21481, the 95^th^ percentile (significant) threshold and 67^th^ percentile (suggestive) threshold were 4.8 and 3.8 LOD, respectively. For trait 21483, the 95^th^ percentile (significant) threshold and 67^th^ percentile (suggestive) threshold were 4.5 and 3.6 LOD, respectively. For trait 21485, the 95^th^ percentile (significant) threshold and 67^th^ percentile (suggestive) threshold were 4.7 and 3.7 LOD, respectively. For trait 21487, the 95^th^ percentile (significant) threshold and 67^th^ percentile (suggestive) threshold were 4.7 and 3.7 LOD, respectively.

QTL scans on small subsets of BXDs, as in our study, are expected only to identify QTLs of large effect. For example, a mapping population comprised of 20 BXD strains with 4 replicates per strain is only powered to detect a QTL explaining 50% of the trait variance, whereas a population size of ~80 strains with 4 replicates per strain is well powered to detect a QTL explaining 20% of the trait variation at 80% power (Andreux et al. 2012; Ashbrook et al. 2021).

### Candidate Gene Search

A 1.5-LOD drop from the peak marker (HK mapping) was used to define an approximate ~95% confidence interval (CI) for a QTL of interest (Alonso-Blanco, Koornneef, and van Ooijen 2006). For each QTL CI, a complete list of mouse reference genes was generated using the UCSC Genome Browser (RRID:SCR_005780) Table Browser Tool (Group: Genes and Gene Predictions; Track: NCBI RefSeq; Genome: GRCm38/mm10).

Genes in each QTL CI were then prioritized based on several criterion. First, all variants distinguishing B6 and D2 within each QTL CI were identified using Sanger’s Mouse Genome Project Mouse_SNPViewer/rel-15050 (RRID:SCR_011784). Specifically, SNPs, InDels, and structural variants (SVs) distinguishing B6 and D2 and their predicted consequences were retrieved and compared (by gene symbol) to reference genes located within the boundaries of each QTL CI. Next, legacy mRNA datasets available in GN for BXD brain tissue were queried to identify genes located in each QTL CI whose expression was modulated by variants located within or near the location of the gene itself—a cis expression QTL (eQTL). Brain region data sets (cortex, hippocampus, hypothalamus, nucleus accumbens, and striatum) were selected based on known functional regulation by cannabinoid receptor signaling, moderate to high expression of CB1R in each region, and functional involvement of each region in initial response traits (*e.g*., motor, hypothermia, and nociception). Datasets included: Hippocampus Consortium M430v2 (Jun06) [GN Accession: GN110 and GEO Series: GSE84767]; INIA Hypothalamus Affy MoGene 1.0 ST (Nov10) [GN Accession: GN281 and GEO Series: GSE36674]; HQF BXD Neocortex ILM6v1.1 (Feb08) RankInv [GN Accession: GN284]; VCU BXD NA Et vs Sal M430 2.0 (Oct07) [GN Accession: GN156]; VCU BXD PFC Sal M430 2.0 (Dec06) RMA [GN Accession: GN135; GEO Series: GSE28515]; HQF Striatum Affy Mouse Exon 1.0ST Gene Level (Dec09) RMA [GN Accession: GN399]. Cis eQTL data are summarized in **Supplemental Table 2**. Probes that overlap polymorphic SNPs and have cis eQTLs with higher expression associated with the B6 (*B*) allele (ILM6v1.1 and M430 data sets only) are flagged in this table as expression measurements that could be biased in the direction of *B* alleles (higher expression relative to D2, or *D*, alleles) due to technical probe hybridization artifacts. Finally, a search for relevant biological function (*i.e*. endocannabinoid/cannabinoid receptor signaling) was conducted using the RatsPub web service (RRID:SCR_018905) to search through PubMed (RRID:SCR_004846) literature abstracts for associations between cannabinoid-related terms and gene symbols for each QTL CI. A secondary search was performed for high priority candidate genes using *RatsPub* to place these genes in a broader context of addiction and psychiatric diseases.

### BXD Exploratory Phenome Analysis

Two recent studies quantified initial responses to multiple drugs of abuse (cocaine, alcohol, and morphine) or intravenous cocaine self-administration among BXDs (Dickson et al. 2016; Philip et al. 2010). Both studies also included large numbers of BXDs, including at least 12 of the strains used to generate our THC initial response data. Associated drug response traits from these studies have been deposited into the GN *BXD Published Phenotypes* dataset. We selected BXD drug response and cocaine self-administration traits from both of these studies for comparison with THC initial response traits generated by our study. Trait data was obtained with the *Get Any* search function in GN using the following input: 2681503 19958391(PubMed IDs). This resulted in 790 traits. The BXD legacy traits mined from GN were retained for correlation analysis if there were at least 12 matched BXD strains shared with our THC initial response traits. This resulted in retention of 757/790 traits. Traits were further filtered if descriptions included the terms “Morphine response”, “morphine withdrawal”, “Cocaine response”, “Intravenous cocaine”, OR “Ethanol response” and DID NOT include the terms “Saline control”, “inactive”, OR “unconditioned”. A final total of 302 traits (excluding the four initial THC response traits) were included for correlation analysis. Traits were assigned to general categories based on phenotype and whether the trait was measured in females, males, or both. Category descriptions along with GN record IDs, trait descriptions, and values for drug response and cocaine self-administration can be found in **Supplemental Table 4**. The Pearson correlation coefficient and corresponding *p*-values were calculated for each of the four THC initial response traits and all 302 BXD legacy drug response traits. Significant correlations were associated with a *p* < 0.05.

## ACKNOWLEDGEMENTS

This research was funded in part by a Cornet Award from the UTHSC Office of Research to Drs. Mulligan, Moore, and Jones and through the UTHSC Center for Integrative and Translational Genetics Mouse Strain and Pilot Projects program. We acknowledge and appreciate the continued support of the NIDA Drug Supply Program for providing THC to Dr. Moore. We also acknowledge talented senior research associates Sufiya Khanam, Christine Watkins, Trevor Houseal, and Tom Shapaker who assisted with phenotyping.

---

**Supplemental Figure 1.**
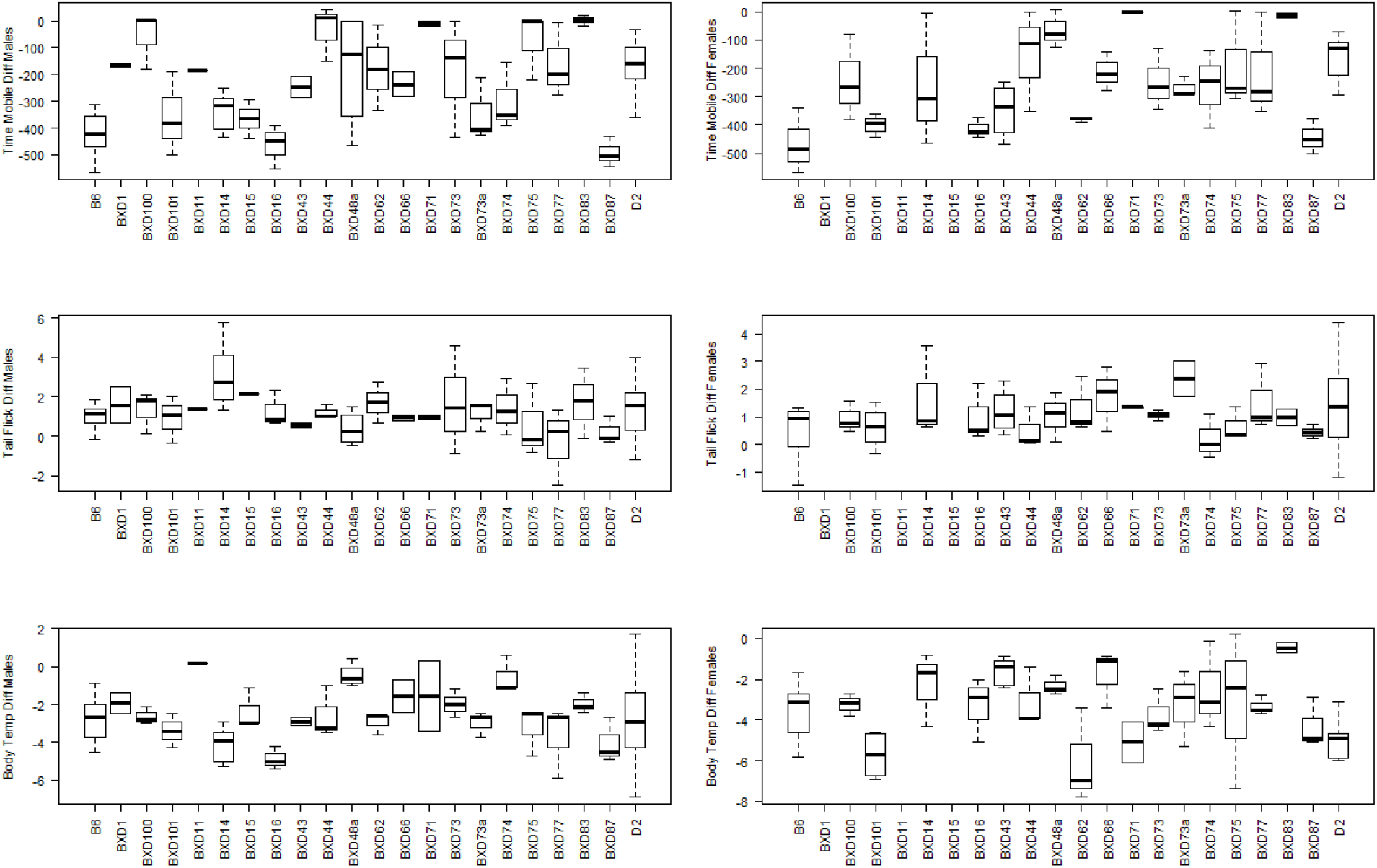
Distribution of initial response traits among B6, D2, and their recombinant inbred BXD progeny. Strains are shown on the x-axis and the y-axis shows initial response to THC treatment (difference between baseline day 0 and day 1). Male and female responses for each trait are shown in the left and right columns, respectively.

**Supplemental Figure 2.**
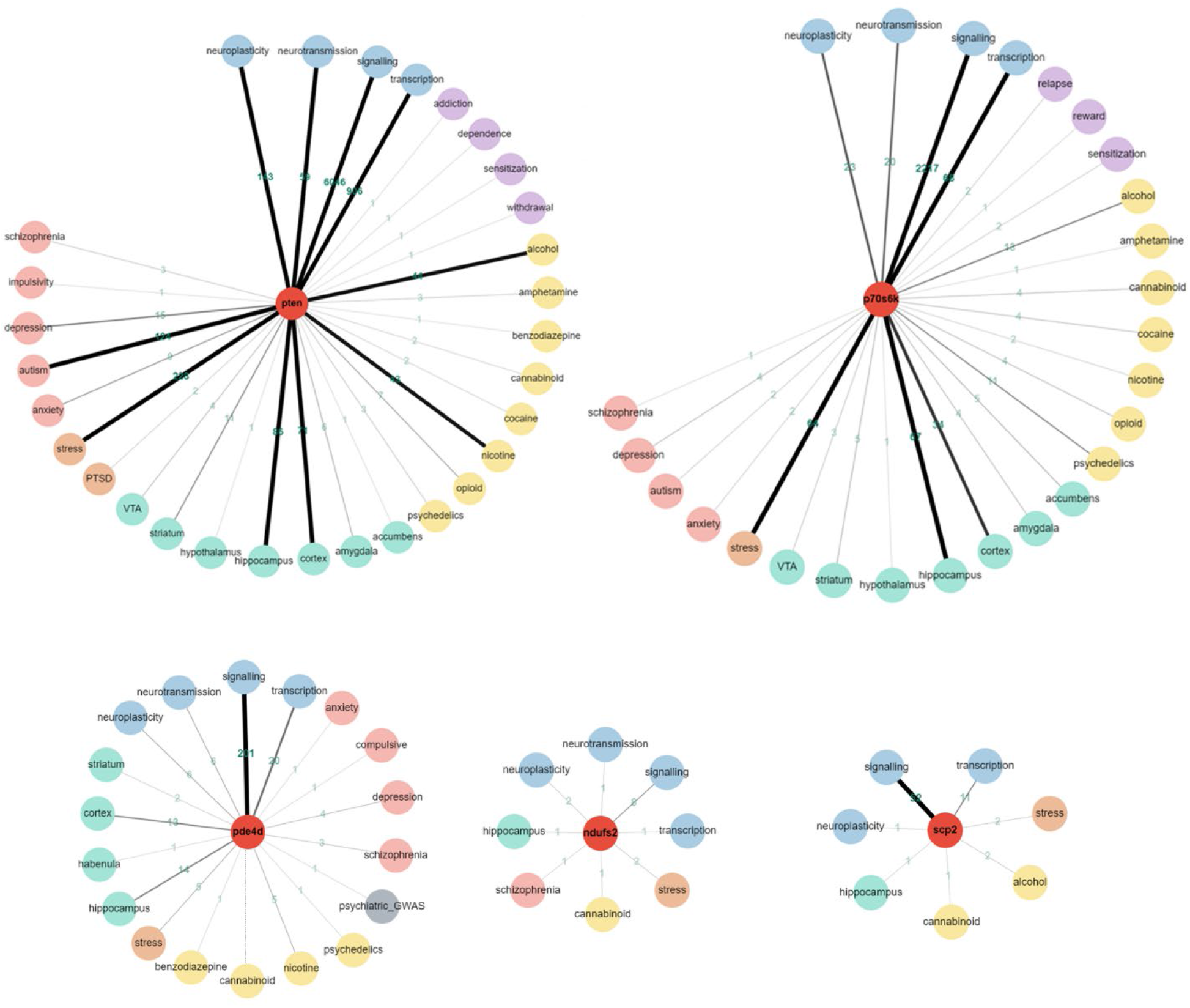
Associations between QTGs and signaling, brain region, addiction, and psychiatric disease terms from the RatsPub Resource. The center node (red) represents the QTG. Terms are shown as nodes (colored by group: blue for signaling terms, yellow for drug terms, purple for addiction terms, green for brain region terms, brown for stress terms and orange for psychiatric terms) radiating around the center node. Edges represent literature associations from pubmed and the edge thickness increases as the number of associations increases. For *Pde4d*, association to the cannabinoid term is indirect (dashed line) and is linked to PDE4 activity and the edge (de Jong et al. 2015) connected to the psychiatric_GWAS term is for an association (*p* = 5E-8) between variants in the human gene (rs159497) and forced expiratory volume in 1 second (occupational environmental exposures interaction).

## REFERENCES

Adermark, L., S. Jonsson, M. Ericson, and B. Soderpalm. 2011. ‘Intermittent ethanol consumption depresses endocannabinoid-signaling in the dorsolateral striatum of rat’, Neuropharmacology, 61: 1160–5.

Alonso-Blanco, C., M. Koornneef, and J. W. van Ooijen. 2006. ‘QTL analysis’, Methods Mol Biol, 323: 79–99.

Alvarez-Jaimes, L., I. Polis, and L. H. Parsons. 2009. ‘Regional Influence of Cannabinoid CB1 Receptors in the Regulation of Ethanol Self-Administration by Wistar Rats’, Open Neuropsychopharmacol J, 2: 77–85.

Alvarez-Jaimes, L., D. G. Stouffer, and L. H. Parsons. 2009. ‘Chronic ethanol treatment potentiates ethanol-induced increases in interstitial nucleus accumbens endocannabinoid levels in rats’, J Neurochem, 111: 37–48.

Andreux, P. A., E. G. Williams, H. Koutnikova, R. H. Houtkooper, M. F. Champy, H. Henry, K. Schoonjans, R. W. Williams, and J. Auwerx. 2012. ‘Systems genetics of metabolism: the use of the BXD murine reference panel for multiscalar integration of traits’, Cell, 150: 1287–99.

Ashbrook, D., D. Arends, P. Prins, M. Mulligan, S. Roy, E. Williams, C. Lutz, A. Valenzuela, C Bohl, J. Ingels, M. McCarty, A. Centeno, R. Hager, J. Auwerx, L. Lu, and R. Williams. 2021. ‘A Platform for Experimental Precision Medicine: The Extended BXD Mouse Family’, Cell Systems, In press.

Ashbrook, D. G., S. Roy, B. G. Clifford, T. Riede, M. L. Scattoni, D. H. Heck, L. Lu, and R. W. Williams. 2018. ‘Born to Cry: A Genetic Dissection of Infant Vocalization’, Front Behav Neurosci, 12: 250.

Boucher, A. A., J. C. Arnold, L. Duffy, P. R. Schofield, J. Micheau, and T. Karl. 2007. ‘Heterozygous neuregulin 1 mice are more sensitive to the behavioural effects of Delta9-tetrahydrocannabinol’, Psychopharmacology (Berl), 192: 325–36.

Boucher, A. A., G. E. Hunt, J. Micheau, X. Huang, I. S. McGregor, T. Karl, and J. C. Arnold. 2011. ‘The schizophrenia susceptibility gene neuregulin 1 modulates tolerance to the effects of cannabinoids’, Int J Neuropsychopharmacol, 14: 631–43.

Caille, S., L. Alvarez-Jaimes, I. Polis, D. G. Stouffer, and L. H. Parsons. 2007. ‘Specific alterations of extracellular endocannabinoid levels in the nucleus accumbens by ethanol, heroin, and cocaine self-administration’, J Neurosci, 27: 3695–702.

Ceccarini, J., C. Casteels, M. Koole, G. Bormans, and K. Van Laere. 2013. ‘Transient changes in the endocannabinoid system after acute and chronic ethanol exposure and abstinence in the rat: a combined PET and microdialysis study’, Eur J Nucl Med Mol Imaging, 40: 1582–94.

Cherry, J. A., and R. L. Davis. 1999. ‘Cyclic AMP phosphodiesterases are localized in regions of the mouse brain associated with reinforcement, movement, and affect’, J Comp Neurol, 407: 287–301.

Clarke, R. B., and L. Adermark. 2010. ‘Acute ethanol treatment prevents endocannabinoid-mediated long-lasting disinhibition of striatal output’, Neuropharmacology, 58: 799–805.

Compton, D. R., M. D. Aceto, J. Lowe, and B. R. Martin. 1996. ‘In vivo characterization of a specific cannabinoid receptor antagonist (SR141716A): inhibition of delta 9-tetrahydrocannabinol-induced responses and apparent agonist activity’, J Pharmacol Exp Ther, 277: 586–94.

de Jong, K., J. M. Vonk, W. Timens, Y. Bosse, D. D. Sin, K. Hao, H. Kromhout, R. Vermeulen, D. S. Postma, and H. M. Boezen. 2015. ‘Genome-wide interaction study of gene-by-occupational exposure and effects on FEV1 levels’, J Allergy Clin Immunol, 136: 1664–72 e14.

Demontis, D., V. M. Rajagopal, T. E. Thorgeirsson, T. D. Als, J. Grove, K. Leppala, D. F. Gudbjartsson, J. Pallesen, C. Hjorthoj, G. W. Reginsson, T. Tyrfingsson, V. Runarsdottir, P. Qvist, J. H. Christensen, J. Bybjerg-Grauholm, M. Baekvad-Hansen, L. M. Huckins, E. A. Stahl, A. Timmermann, E. Agerbo, D. M. Hougaard, T. Werge, O. Mors, P. B. Mortensen, M. Nordentoft, M. J. Daly, H. Stefansson, K. Stefansson, M. Nyegaard, and A. D. Borglum. 2019. ‘Genome-wide association study implicates CHRNA2 in cannabis use disorder’, Nat Neurosci, 22: 1066–74.

Dickson, P. E., M. M. Miller, M. A. Calton, J. A. Bubier, M. N. Cook, D. Goldowitz, E. J. Chesler, and G. Mittleman. 2016. ‘Systems genetics of intravenous cocaine self-administration in the BXD recombinant inbred mouse panel’, Psychopharmacology (Berl), 233: 701–14.

Ellgren, M., S. M. Spano, and Y. L. Hurd. 2007. ‘Adolescent cannabis exposure alters opiate intake and opioid limbic neuronal populations in adult rats’, Neuropsychopharmacology, 32: 607–15.

Fourgeaud, L., S. Mato, D. Bouchet, A. Hemar, P. F. Worley, and O. J. Manzoni. 2004. ‘A single in vivo exposure to cocaine abolishes endocannabinoid-mediated long-term depression in the nucleus accumbens’, J Neurosci, 24: 6939–45.

Grueter, B. A., H. B. Gosnell, C. M. Olsen, N. L. Schramm-Sapyta, T. Nekrasova, G. E. Landreth, and D. G. Winder. 2006. ‘Extracellular-signal regulated kinase 1-dependent metabotropic glutamate receptor 5-induced long-term depression in the bed nucleus of the stria terminalis is disrupted by cocaine administration’, J Neurosci, 26: 3210–9.

Guegan, T., J. P. Cebria, R. Maldonado, and M. Martin. 2016. ‘Morphine-induced locomotor sensitization produces structural plasticity in the mesocorticolimbic system dependent on CB1-R activity’, Addict Biol, 21: 1113–26.

Han, S., B. Z. Yang, H. R. Kranzler, D. Oslin, R. Anton, L. A. Farrer, and J. Gelernter. 2012. ‘Linkage analysis followed by association show NRG1 associated with cannabis dependence in African Americans’, Biol Psychiatry, 72: 637–44.

Harenza, J. L., P. P. Muldoon, M. De Biasi, M. I. Damaj, and M. F. Miles. 2014. ‘Genetic variation within the Chrna7 gene modulates nicotine reward-like phenotypes in mice’, Genes Brain Behav, 13: 213–25.

Hebert-Chatelain, E., T. Desprez, R. Serrat, L. Bellocchio, E. Soria-Gomez, A. Busquets-Garcia, A. C. Pagano Zottola, A. Delamarre, A. Cannich, P. Vincent, M. Varilh, L. M. Robin, G. Terral, M. D. Garcia-Fernandez, M. Colavita, W. Mazier, F. Drago, N. Puente, L. Reguero, I. Elezgarai, J. W. Dupuy, D. Cota, M. L. Lopez-Rodriguez, G. Barreda-Gomez, F. Massa, P. Grandes, G. Benard, and G. Marsicano. 2016. ‘A cannabinoid link between mitochondria and memory’, Nature, 539: 555–59.

Hill, W., and T. Mackay. 2004. ‘D.S. Falconer and Introduction to Quantitative Genetics’, Genetics, 167: 1529–36.

Hitzemann, R., B. Hitzemann, S. Rivera, J. Gatley, P. Thanos, L. L. Shou, and R. W. Williams. 2003. ‘Dopamine D2 receptor binding, Drd2 expression and the number of dopamine neurons in the BXD recombinant inbred series: genetic relationships to alcohol and other drug associated phenotypes’, Alcohol Clin Exp Res, 27: 1–11.

Hryhorowicz, S., M. Walczak, O. Zakerska-Banaszak, R. Slomski, and M. Skrzypczak-Zielinska. 2018. ‘Pharmacogenetics of Cannabinoids’, Eur J Drug Metab Pharmacokinet, 43: 1–12.

Huestis, M. A., D. A. Gorelick, S. J. Heishman, K. L. Preston, R. A. Nelson, E. T. Moolchan, and R. A. Frank. 2001. ‘Blockade of effects of smoked marijuana by the CB1-selective cannabinoid receptor antagonist SR141716’, Arch Gen Psychiatry, 58: 322–8.

Jackson, K. J., X. Chen, M. F. Miles, J. Harenza, and M. I. Damaj. 2011. ‘The neuropeptide galanin and variants in the GalR1 gene are associated with nicotine dependence’, Neuropsychopharmacology, 36: 2339–48.

Ji, S. P., Y. Zhang, J. Van Cleemput, W. Jiang, M. Liao, L. Li, Q. Wan, J. R. Backstrom, and X. Zhang. 2006. ‘Disruption of PTEN coupling with 5-HT2C receptors suppresses behavioral responses induced by drugs of abuse’, Nat Med, 12: 324–9.

Knoll, A. T., K. Jiang, and P. Levitt. 2018. ‘Quantitative trait locus mapping and analysis of heritable variation in affiliative social behavior and co-occurring traits’, Genes Brain Behav, 17: e12431.

Lariviere, W. R., and J. S. Mogil. 2010. ‘The genetics of pain and analgesia in laboratory animals’, Methods Mol Biol, 617: 261–78.

Laughlin, R. E., T. L. Grant, R. W. Williams, and J. D. Jentsch. 2011. ‘Genetic dissection of behavioral flexibility: reversal learning in mice’, Biol Psychiatry, 69: 1109–16.

Ledent, C., O. Valverde, G. Cossu, F. Petitet, J. F. Aubert, F. Beslot, G. A. Bohme, A. Imperato, T. Pedrazzini, B. P. Roques, G. Vassart, W. Fratta, and M. Parmentier. 1999. ‘Unresponsiveness to cannabinoids and reduced addictive effects of opiates in CB1 receptor knockout mice’, Science, 283: 401–4.

Liedhegner, E. S., C. D. Vogt, D. S. Sem, C. W. Cunningham, and C. J. Hillard. 2014. ‘Sterol carrier protein-2: binding protein for endocannabinoids’, Mol Neurobiol, 50: 149–58.

Liu, Q. S., L. Pu, and M. M. Poo. 2005. ‘Repeated cocaine exposure in vivo facilitates LTP induction in midbrain dopamine neurons’, Nature, 437: 1027–31.

Loos, M., T. Mueller, Y. Gouwenberg, R. Wijnands, R. J. van der Loo, Bsik Mouse Phenomics Consortium Neuro, C. Birchmeier, A. B. Smit, and S. Spijker. 2014. ‘Neuregulin-3 in the mouse medial prefrontal cortex regulates impulsive action’, Biol Psychiatry, 76: 648–55.

Maillet, J. C., Y. Zhang, X. Li, and X. Zhang. 2008. ‘PTEN-5-HT2C coupling: a new target for treating drug addiction’, Prog Brain Res, 172: 407–20.

Martin, G. G., D. R. Seeger, A. L. McIntosh, S. Milligan, S. Chung, D. Landrock, L. J. Dangott, M. Y. Golovko, E. J. Murphy, A. B. Kier, and F. Schroeder. 2019. ‘Sterol Carrier Protein-2/Sterol Carrier Protein-x/Fatty Acid Binding Protein-1 Ablation Impacts Response of Brain Endocannabinoid to High-Fat Diet’, Lipids, 54: 583–601.

Monory, K., H. Blaudzun, F. Massa, N. Kaiser, T. Lemberger, G. Schutz, C. T. Wotjak, B. Lutz, and G. Marsicano. 2007. ‘Genetic dissection of behavioural and autonomic effects of Delta(9)-tetrahydrocannabinol in mice’, PLoS Biol, 5: e269.

Mulligan, M. K., K. Mozhui, P. Prins, and R. W. Williams. 2017. ‘GeneNetwork: A Toolbox for Systems Genetics’, Methods Mol Biol, 1488: 75–120.

Mulligan, M. K., W. Zhao, M. Dickerson, D. Arends, P. Prins, S. A. Cavigelli, E. Terenina, P. Mormede, L. Lu, and B. C. Jones. 2018. ‘Genetic Contribution to Initial and Progressive Alcohol Intake Among Recombinant Inbred Strains of Mice’, Front Genet, 9: 370.

Muschamp, J. W., and W. A. Carlezon, Jr. 2013. ‘Roles of nucleus accumbens CREB and dynorphin in dysregulation of motivation’, Cold Spring Harb Perspect Med, 3: a012005.

Nestler, E. J. 2016. ‘Reflections on: “A general role for adaptations in G-Proteins and the cyclic AMP system in mediating the chronic actions of morphine and cocaine on neuronal function”‘, Brain Res, 1645: 71–4.

Neuner, S. M., B. P. Garfinkel, L. A. Wilmott, B. M. Ignatowska-Jankowska, A. Citri, J. Orly, L. Lu, R. W. Overall, M. K. Mulligan, G. Kempermann, R. W. Williams, K. M. O’Connell, and C. C. Kaczorowski. 2016. ‘Systems genetics identifies Hp1bp3 as a novel modulator of cognitive aging’, Neurobiol Aging, 46: 58–67.

Newbury, A. J., and G. D. Rosen. 2012. ‘Genetic, morphometric, and behavioral factors linked to the midsagittal area of the corpus callosum’, Front Genet, 3: 91.

Newell, K. A., T. Karl, and X. F. Huang. 2013. ‘A neuregulin 1 transmembrane domain mutation causes imbalanced glutamatergic and dopaminergic receptor expression in mice’, Neuroscience, 248: 670–80.

Ning, K., L. Pei, M. Liao, B. Liu, Y. Zhang, W. Jiang, J. G. Mielke, L. Li, Y. Chen, Y. H. El-Hayek, M. G. Fehlings, X. Zhang, F. Liu, J. Eubanks, and Q. Wan. 2004. ‘Dual neuroprotective signaling mediated by downregulating two distinct phosphatase activities of PTEN’, J Neurosci, 24: 4052–60.

Olsen, C. M., and Q. S. Liu. 2016. ‘Phosphodiesterase 4 inhibitors and drugs of abuse: current knowledge and therapeutic opportunities’, Front Biol (Beijing), 11: 376–86.

Pan, B., C. J. Hillard, and Q. S. Liu. 2008. ‘Endocannabinoid signaling mediates cocaine-induced inhibitory synaptic plasticity in midbrain dopamine neurons’, J Neurosci, 28: 1385–97.

Panagis, G., B. Mackey, and S. Vlachou. 2014. ‘Cannabinoid Regulation of Brain Reward Processing with an Emphasis on the Role of CB1 Receptors: A Step Back into the Future’, Front Psychiatry, 5: 92.

Parker, C. C., P. E. Dickson, V. M. Philip, M. Thomas, and E. J. Chesler. 2017. ‘Systems Genetic Analysis in GeneNetwork.org’, Curr Protoc Neurosci, 79: 8 39 1–8 39 20.

Parks, C., F. Giorgianni, B. C. Jones, S. Beranova-Giorgianni, B. M. Moore Ii, and M. K. Mulligan. 2019. ‘Comparison and Functional Genetic Analysis of Striatal Protein Expression Among Diverse Inbred Mouse Strains’, Front Mol Neurosci, 12: 128.

Parks, C., B. C. Jones, B. Moore, and M. Mulligan. 2020. ‘Sex and Strain Variation in Initial Sensitivity and Rapid Tolerance to D9–Tetrahydrocannabinol’, Cannabis and Cannabinoid Research, In Press.

Parsons, L. H., and Y. L. Hurd. 2015. ‘Endocannabinoid signalling in reward and addiction’, Nat Rev Neurosci, 16: 579–94.

Pasman, J. A., K. J. H. Verweij, Z. Gerring, S. Stringer, S. Sanchez-Roige, J. L. Treur, A. Abdellaoui, M. G. Nivard, B. M. L. Baselmans, J. S. Ong, H. F. Ip, M. D. van der Zee, M. Bartels, F. R. Day, P. Fontanillas, S. L. Elson, Team andMe Research, H. de Wit, L. K. Davis, J. MacKillop, Consortium Substance Use Disorders Working Group of the Psychiatric Genomics, Consortium International Cannabis, J. L. Derringer, S. J. T. Branje, C. A. Hartman, A. C. Heath, P. A. C. van Lier, P. A. F. Madden, R. Magi, W. Meeus, G. W. Montgomery, A. J. Oldehinkel, Z. Pausova, J. A. Ramos-Quiroga, T. Paus, M. Ribases, J. Kaprio, M. P. M. Boks, J. T. Bell, T. D. Spector, J. Gelernter, D. I. Boomsma, N. G. Martin, S. MacGregor, J. R. B. Perry, A. A. Palmer, D. Posthuma, M. R. Munafo, N. A. Gillespie, E. M. Derks, and J. M. Vink. 2018. ‘GWAS of lifetime cannabis use reveals new risk loci, genetic overlap with psychiatric traits, and a causal influence of schizophrenia’, Nat Neurosci, 21: 1161–70.

Perez-Torres, S., X. Miro, J. M. Palacios, R. Cortes, P. Puigdomenech, and G. Mengod. 2000. ‘Phosphodiesterase type 4 isozymes expression in human brain examined by in situ hybridization histochemistry and[3H]rolipram binding autoradiography. Comparison with monkey and rat brain’, J Chem Neuroanat, 20: 349–74.

Philip, V. M., S. Duvvuru, B. Gomero, T. A. Ansah, C. D. Blaha, M. N. Cook, K. M. Hamre, W. R. Lariviere, D. B. Matthews, G. Mittleman, D. Goldowitz, and E. J. Chesler. 2010. ‘High-throughput behavioral phenotyping in the expanded panel of BXD recombinant inbred strains’, Genes Brain Behav, 9: 129–59.

Phillips, T. J., J. K. Belknap, and J. C. Crabbe. 1991. ‘Use of recombinant inbred strains to assess vulnerability to drug abuse at the genetic level’, J Addict Dis, 10: 73–87.

Puighermanal, E., A. Busquets-Garcia, M. Gomis-Gonzalez, G. Marsicano, R. Maldonado, and A. Ozaita. 2013. ‘Dissociation of the pharmacological effects of THC by mTOR blockade’, Neuropsychopharmacology, 38: 1334–43.

Puighermanal, E., G. Marsicano, A. Busquets-Garcia, B. Lutz, R. Maldonado, and A. Ozaita. 2009. ‘Cannabinoid modulation of hippocampal long-term memory is mediated by mTOR signaling’, Nat Neurosci, 12: 1152–8.

Rinaldi-Carmona, M., F. Barth, M. Heaulme, D. Shire, B. Calandra, C. Congy, S. Martinez, J. Maruani, G. Neliat, D. Caput, and et al. 1994. ‘SR141716A, a potent and selective antagonist of the brain cannabinoid receptor’, FEBS Lett, 350: 240–4.

Rodriguez, L. A., R. Plomin, D. A. Blizard, B. C. Jones, and G. E. McClearn. 1994. ‘Alcohol acceptance, preference, and sensitivity in mice. I. Quantitative genetic analysis using BXD recombinant inbred strains’, Alcohol Clin Exp Res, 18: 1416–22.

Rodriguez, L. A., R. Plomin, D. A. Blizard, B. C. Jones, and G. E. McClearn.1995. ‘Alcohol acceptance, preference, and sensitivity in mice. II. Quantitative trait loci mapping analysis using BXD recombinant inbred strains’, Alcohol Clin Exp Res, 19: 367–73.

Sachse-Seeboth, C., J. Pfeil, D. Sehrt, I. Meineke, M. Tzvetkov, E. Bruns, W. Poser, S. V. Vormfelde, and J. Brockmoller. 2009. ‘Interindividual variation in the pharmacokinetics of Delta9-tetrahydrocannabinol as related to genetic polymorphisms in CYP2C9’, Clin Pharmacol Ther, 85: 273–6.

Serrano, A., and L. H. Parsons. 2011. ‘Endocannabinoid influence in drug reinforcement, dependence and addiction-related behaviors’, Pharmacol Ther, 132: 215–41.

Sherva, R., Q. Wang, H. Kranzler, H. Zhao, R. Koesterer, A. Herman, L. A. Farrer, and J. Gelernter. 2016. ‘Genome-wide Association Study of Cannabis Dependence Severity, Novel Risk Variants, and Shared Genetic Risks’, JAMA Psychiatry, 73: 472–80.

Stringer, S., C. C. Minica, K. J. Verweij, H. Mbarek, M. Bernard, J. Derringer, K. R. van Eijk, J. D. Isen, A. Loukola, D. F. Maciejewski, E. Mihailov, P. J. van der Most, C. Sanchez-Mora, L. Roos, R. Sherva, R. Walters, J. J. Ware, A. Abdellaoui, T. B. Bigdeli, S. J. Branje, S. A. Brown, M. Bruinenberg, M. Casas, T. Esko, I. Garcia-Martinez, S. D. Gordon, J. M. Harris, C. A. Hartman, A. K. Henders, A. C. Heath, I. B. Hickie, M. Hickman, C. J. Hopfer, J. J. Hottenga, A. C. Huizink, D. E. Irons, R. S. Kahn, T. Korhonen, H. R. Kranzler, K. Krauter, P. A. van Lier, G. H. Lubke, P. A. Madden, R. Magi, M. K. McGue, S. E. Medland, W. H. Meeus, M. B. Miller, G. W. Montgomery, M. G. Nivard, I. M. Nolte, A. J. Oldehinkel, Z. Pausova, B. Qaiser, L. Quaye, J. A. Ramos-Quiroga, V. Richarte, R. J. Rose, J. Shin, M. C. Stallings, A. I. Stiby, T. L. Wall, M. J. Wright, H. M. Koot, T. Paus, J. K. Hewitt, M. Ribases, J. Kaprio, M. P. Boks, H. Snieder, T. Spector, M. R. Munafo, A. Metspalu, J. Gelernter, D. I. Boomsma, W. G. Iacono, N. G. Martin, N. A. Gillespie, E. M. Derks, and J. M. Vink. 2016. ‘Genome-wide association study of lifetime cannabis use based on a large meta-analytic sample of 32 330 subjects from the International Cannabis Consortium’, Transl Psychiatry, 6: e769.

Zhao, X., L. Yao, F. Wang, H. Zhang, and L. Wu. 2017. ‘Cannabinoid 1 receptor blockade in the dorsal hippocampus prevents the reinstatement but not acquisition of morphine-induced conditioned place preference in rats’, Neuroreport, 28: 565–70.

Zhou, X., and M. Stephens. 2012. ‘Genome-wide efficient mixed-model analysis for association studies’, Nat Genet, 44: 821–4.

Zimmer, A., A. M. Zimmer, A. G. Hohmann, M. Herkenham, and T. I. Bonner. 1999. ‘Increased mortality, hypoactivity, and hypoalgesia in cannabinoid CB1 receptor knockout mice’, Proc Natl Acad Sci U S A, 96: 5780–5.

